# Fine–mapping identifies *NAD–ME1* as a candidate underlying a major locus controlling temporal variation in primary and specialized metabolism in Arabidopsis

**DOI:** 10.1101/2020.09.07.285429

**Authors:** Marta Francisco, Daniel J. Kliebenstein, Víctor M. Rodríguez, Pilar Soengas, Rosaura Abilleira, María E. Cartea

## Abstract

Plant metabolism is modulated by a complex interplay between internal signals and external cues. A major goal of all quantitative metabolomic studies is to clone the underlying genes to understand the mechanistic basis of this variation. Using fine-scale genetic mapping, in this work we report the identification and initial characterization of *NAD-DEPENDENT MALIC ENZYME 1* (*NAD*-*ME1*) as the candidate gene underlying the pleiotropic network Met.II.15 QTL controlling variation in plant metabolism and circadian clock outputs in the Bay × Sha Arabidopsis population. Transcript abundance and promoter analysis in *NAD-ME1*^Bay-0^ and *NAD-ME1*^Sha^ alleles confirmed allele-specific expression that appears to be due a polymorphism disrupting a putative circadian cis-element binding site. Analysis of T-DNA insertion lines and heterogeneous inbred families (HIFs) showed that transcript variation of the *NAD-ME1* gene led to temporal shifts of tricarboxylic acid cycle (TCA) intermediates, glucosinolate (GSL) accumulation and altered regulation of several GSL biosynthesis pathway genes. Untargeted metabolomics analyses reveal complex regulatory networks of *NAD-ME1* dependent upon the day-time. The mutant lead to shifts in plant primary metabolites, cell-wall components, isoprenoids, fatty acids and plant immunity phytochemicals, among others. Our findings suggest that *NAD-ME1* may act as a key component to coordinate plant primary and secondary metabolism in a time-dependent manner.

## Introduction

Plants produce large arsenals of structurally and diverse metabolic pathways with metabolites being generally categorized as primary metabolites or secondary metabolites (*1*). Primary metabolites, including those of the tricarboxylic acid cycle (TCA), sugars, amino acids and lipids are involved in central processes of growth and development producing all the necessary building blocks for cells and the resulting biomass. Secondary metabolites such as phenolics, terpenoids, alkaloids, and glucosinolates (GSLs) are specialized compounds that play critical roles in plant adaptation under stressful environmental events (*2, 3*). Given the central importance of metabolism in any physiological or developmental process, metabolic networks must be fine-tuned to make the most efficient use of available resources. The plant metabolome, however, is not a stable entity. Fluxes and metabolite concentrations are variables while enzyme activities are under genetic control (*4*). The precise coordination of metabolic fluxes in response to diverse endogenous signals and environmental conditions has led to strong interest in understanding how plant metabolism is regulated (*5*).

One of the key components in integrating environmental signals and regulating metabolite levels is the circadian clock (*6*). This coordination is critical for local adaptation because it allows plants to anticipate predictable daily environmental changes like sunrise and sunset and adjust development and physiology (*7*). In the model plant, Arabidopsis, several works revealed that transcript amounts of key enzymes from central metabolism are regulated by the circadian clock (*8, 9*). This regulation coordinates primary metabolite levels, reserve turnover, and growth with the daily cycle of light and darkness (*10–15*). In addition, evidence that specialized metabolites levels can oscillate rhythmically during a day is beginning to accumulate (*16–20*). An example of this are the GSLs, a major class of sulphur-containing secondary metabolites present in the members of family *Brassicaceae* that provide resistance against pathogens and insects (*21*). These compounds show a circadian pattern of accumulation in Arabidopsis and in crops from the same family (*20, 22, 23*). This suggests a complex and intricate biological timing mechanism that governs metabolism daily behavior. Thus, unraveling the genetic basis of this variation can remarkably enhance our understanding of plant integral regulatory systems for metabolic traits (*24*).

Genetic connections between both primary and secondary metabolism and circadian outputs begin to be elucidated in Arabidopsis. The study of metabolomic variation in the Bayreuth-0 (Bay-0, CS954) × Shahdara (Sha, CS929) recombinant inbred line (RIL) population identified a number of genomic regions associated with metabolite variation, namely metabolic quantitative trait loci (Met.QTL) (*25, 26*). Further transcriptomic analysis of the same RIL population revealed that several QTLs for natural variation altering primary metabolism were linked to the expression of the circadian clock output networks (Kerwin *et al*., 2011). Interestingly, seven of those QTL clusters (Met.chromosome.centimorgan [cM]), Met.II.15, Met.II.47, Met.III.04, Met.IV.65, Met.V.67, AOP and ELONG found for the circadian clock and metabolomics collocated with previously identified cis-eQTL (natural variation in gene expression that map to their physical position) hot spots for GSL pathway related transcripts (West *et al*., 2007; Wentzell *et al*., 2007). Molecular cloning of major metabolic/clock QTLs are beginning to show that metabolomics variation may often identify key physiological regulators that are important control nodes for the metabolome with pleiotropic effects across plant circadian rhythms. In support of this, the AOP and ELONG loci play a crucial role in GSL biosynthesis determining aliphatic GSL accumulation and structure but also, AOP is involved in control of onset of flowering and circadian period alteration (*27, 28*). Similarly, Met. II.47 QTL caused by natural variation between the Bay and Sha alleles at *EARLY FLOWERING3* (*ELF3*), has been linked to altering flowering time, shade-avoidance, hypocotyl elongation, circadian clock oscillation and metabolic variation (*29–31*). Besides, the Met.V.67 locus, controlled by natural variation in *METABOLIC NETWORK MODULATOR 1* (*MNM1*), affected primary metabolism accumulation within Arabidopsis in a time-dependent fashion (*32*). Thus, cloned genes are illuminating new and unexpected mechanistic aspects of plant biology.

To provide new insights into the modular behavior of metabolic processes and their regulation, the objective of the present work was to identify and clone candidate genes in the genomic region of the pleiotropic network Met.II.15 QTL controlling variation in plant metabolism and circadian clock outputs within Arabidopsis. To accomplish this, we utilized a combination of fine*–*scale genetic mapping, using heterogeneous inbred families (HIFs), and *in silico* candidate gene identification. These analyses detected *NAD-DEPENDENT MALIC ENZYME 1* (*NAD–ME1*) as the most likely candidate gene underlying the Met.II.15 QTL. Transcript abundance and promoter analysis in *NAD–ME1*^Bay-0^ and *NAD–ME1*^Sha^ alleles confirmed allele-specific expression that appears to be due a polymorphism disrupting a putative circadian cis-element binding site. Analysis of T–DNA insertion lines and HIFs showed that transcripts variation of the *NAD–ME1* gene led to temporal shifts of tricarboxylic acid (TCA) intermediates, GSL accumulation and altered regulation of several GSL biosynthesis pathway genes. Untargeted metabolomic analyses reveal complex regulatory networks of *NAD*–*ME1* dependent upon the day–time. The mutant lead to shifts in plant primary metabolites, cell–wall components, isoprenoids, fatty acids and plant immunity phytochemicals, among others. Together, these data show that *NAD–ME1* may act as a key component to coordinate plant primary and secondary metabolism in a time dependent manner.

## Materials and Methods

### Mapping population and fine–mapping strategy

To confirm the effects of the locus in an isogenic background Near Isogenic Lines (NILs) were developed using a HIF following the idea published by Tuintra *et al*. (*33*). This powerful approach uses the residual heterozygosity in early generations of RILs. The Bay-0 × Sha RIL population at F6 showed approximately 97% homozygosity in each line (420 lines in total). This resulted in the presence of residual heterozygosity in at least a single RIL at almost all genome positions. In the present study homozygous NILs were created from the progeny of the line RIL364 (from the Bay-0 × Sha population) that show a single residual heterozygous region around marker MSAT2.28 where Met.II.15 QTL was originally mapped (*25–27, 33*). Seeds of RIL364 at F6 stage were kindly provided by Dr. Olivier Loudet (INRA, Versailles, France). For fine mapping, a homozygous RIL364 [Bay-0] and RIL364 [Sha] pair was crossed to recreate heterozygosity (HIF364) at Met.II.15 QTL in an isogenic background and the F1 selfed to make F2. PCR-based markers flanking Met.II.15 were developed and used to screen 1,600 F2 progeny to isolate individuals fixed for the parental alleles in the interval as well as possible recombinants within the heterozygous region between markers MSAT200897, MSAT2.28 and IND268. All recombinant progeny were then self–fertilized to generate F3 seeds to validate the F2 measurements. To more precisely determine the recombination breakpoints we used data sequence from 1001 Genomes project (*34*) to computationally predict INDELs between Bay-0 and Sha. With this approach, additional markers between markers MSAT2.28 and IND268 were developed and used to genotype all recombinants. Primers were designed using software Primer 3.0 (http://bioinfo.ut.ee/primer3-0.4.0/). DNA was extracted using the Promega DNA Purification System. PCR products were run in 2% agarose gel stained with EtBr and the genotype of the plant was assessed. The primers used for the PCRs are listed in Table S1.

### T–DNA insertion lines

The T–DNA insertion line Sail–374–A02 (At2g13560), was kindly provided by Dr. Tronconi (Universidad Nacional de Rosario, Rosario, Argentina). Additional T–DNA insertion lines SALK_024285 (AT2g13540), SALK_090811 (At2g13560) and SALK_126034 (At2g13610) were obtained from the Nottingham Arabidopsis Stock Center (http://www.arabidopsis.info/). Lines with homozygous insertions were identified by PCR–based genotyping and corroborated by RT–qPCR. *NAD–ME1* mutant lines do not showed altered expression of the neighboring At2g13550 gene. Either *NAD–ME1* mutants not showed visual phenotypic alterations under normal growth conditions (Fig. S1). Since both *NAD–ME1* mutants tested displayed similar phenotypes, results presented here are based on Sail-374-A02 (*NAD–ME1*) mutant. Analysis of variance (ANOVA) was used to compare GSL variation between T–DNA lines from the three candidate genes (At2g13560, At2g13540 and AT2g13610) and Col-0 (wild–type). Each genotype account with 24 independent measurements conducted across two independent experiments.

### Plant growth and harvest conditions

All seeds were planted in potting soil stratified at 4°C in the dark for 3 days to optimize germination using 3.80 × 3.80 cm pots. For fine–mapping analysis, all plants from the appropriate genotypes were grown in long-day (LD) conditions (16/8h light/dark) photoperiods in a growth chamber under controlled conditions (100–120 μE light intensity and 22°C of temperature and 60% of humidity). Rosette leaves were used for all experiments. To test different *NAD–ME1* allele’s effects upon cycling within the GSL network the appropriate genotypes were tested under LD photoperiod and harvested after three weeks of germination. In all the experiments, plants were sown in a randomized block design with two complete independent replicates of the whole experiment. Sampling was performed at various Zeitgeber times (ZT), starting just before dawn (ZT0). To study the effect of *NAD–ME1* alleles on transcript levels of GSL related genes as well as for metabolomics analysis, samples were harvested at ZT4 and ZT8. In each experiment, three individual plants per genotype were harvested at each time point for sampling. This provides six total samples per genotype per time point across the whole dataset. For harvesting, leaves were cut placed in plastic scintillation vials and immediately frozen in liquid nitrogen. During the dark phase, the plants were sampled in the presence of low–intensity green lamp. Plant material was stored at −80°C.

### Real time PCR

Total RNA was extracted using the RNeasy kit (QIAGEN, Germany). To remove any traces of genomic DNA, the RNA was treated with DNase by following the manufacturer instructions. Samples of RNA were reverse transcribed using the GoScript™ reverse transcription system and oligo (dT20) (Promega). RT–qPCR was performed in a 20 μl reaction with the Fast Start Universe SYBR Green Master (ROX) mix (Roche, IN, USA). The expression levels were normalized to glyceraldehyde-3-phosphate-dehydrogenase (GADPH). RT-qPCRs were carried out on a 7500 Real Time PCR System (Applied Biosystem, Forster City, CA, USA) and primer efficiency was calculated using the LingRegPCR software (*35*). Statistical significance was calculated using a Student’s t test to compare the relative gene expression among genotypes. All the primer pairs used are listed in the Table S1.

### GSL identification and quantification

Sample extraction and desulfation, were performed according to Kliebenstein *et al*. (*36*) with minor modifications. Ten microliters of the desulfo-GLS extract were used to identify and quantify the GSLs. The chromatographic analyses were carried out on an Ultra-High-Performance Liquid-Chromatograph (UHPLC Nexera LC-30AD; Shimadzu) equipped with a Nexera SIL-30AC injector and one SPD-M20A UV/VIS photodiode array detector. The UHPLC column was aXSelect HSS T3 XP ColumnC18 protected with a C18 guard cartridge. The oven temperature was set at 30°C. Compounds were separated using the following method in aqueous acetonitrile, with a flow of 0.5 mL min^−1^: 1.5 min at 100% H_2_O, an 11 min gradient from 0% to 25% (v/v) acetonitrile, 1.5 min at 25% (v/v) acetonitrile, a minute gradient from 25% to 0% (v/v) acetonitrile, and a final 3 min at 100% H_2_O. Data was recorded on a computer with the LabSolutions software (Shimadzu). All GSLs were quantified at 229 nm by using glucotropaeolin (GTP, monohydrate from Phytoplan, Diehm & Neuberger GmbH, Heidelberg, Germany) as internal standard and quantified by comparison to purified standards. We reported the concentration (μmol cm^2^ of FW) of individual GSL compounds as well as the sums of total aliphatic and indolic GSLs, two classes of GSLs based on the amino acid from which they have derived. ANOVA was used to compare individual and total GSL variation between Bay-0 and Sha alleles of each molecular marker at Met.II.15 QTL region. Multiple comparisons comparing GSL traits between genotypes were made post-hoc using Tukey's t-test with P ≤ 0.05 within the model.

### Co-expression network analysis

Co-expression data for each studied gene was obtained from ATTED-II version c4.1 that was calculated including 1388 microarray experiments (*37*). To filter candidate genes in the final interval QTL we search for significant correlations between expression variation of candidate genes, GSL-related genes and circadian core clock genes. To construct the network, we filtered the first 30 genes correlated with *NAD–ME1*. Among them, we selected only the genes with a cis–eQTL in the Bay-0 × Sha RIL population (*26*). The selected genes were used as bait genes. Then, co–expression relationship between the bait genes and their directly–connected genes were used to construct a network visualized using NetworkDrawer (*38*). Genes were subsequently interpreted by functional enrichment analysis based on Gene Ontology (GO) to identify and rank overrepresented functional categories in the network.

### Metabolomic analysis

For metabolomic composition analysis we used an ultra–performance liquid chromatography coupled with electrospray ionization quadrupole (Thermo Dionex Ultimate 3000 LC) time–of–flight mass spectrometry (UPLC–Q–TOF–MS/MS) (Bruker Compact™) with a heated electrospray ionization (ESI) source. Chromatographic separation was performed in a Kinetex™ C18 LC Column (2.1□×□100◻mm 1.7□μm pore size) using a binary gradient solvent mode consisting of 0.1% formic acid in water (solvent A) and acetonitrile (solvent B). The following gradient was used: 3 % B (0-4 min), from 3% to 25 % B (4-16 min), from 25 to 80% B (16-25min), from 80 to 100% B (25-30 min), hold 100% B until 32 min, from 100% to 3% B (32-33 min), hold 3% B until 36 min. The injection volume was 5 μL, the flow rate was established at 0.4 ml/min and column temperature was controlled at 35 °C. MS analysis was operated in spectra acquisition range from 50 to 1200 m/z. Both polarities (±) of ESI mode were used under the following specific conditions: gas flow 9 l/min, nebulizer pressure 38 psi, dry gas 9 l/min, and dry temperature 220 °C. Capillary and end plate offset were set to 4500 and 500 V, respectively. MS/MS analysis was performed based on the previously determined accurate mass and RT and fragmented by using different collision energy ramps to cover a range from 15 to 50 eV. The algorithm T–Rex 3D from the MetaboScape 4.0 software (Bruker Daltoniks, Germany) was used for peak alignment and detection. The generated dataset was imported into Metaboanalyst (*39*) to perform statistical analyses. In order to remove non–informative variables, data was filtered using the interquantile range filter (IQR). Moreover, Pareto variance scaling was used to remove the offsets and adjust the importance of high– and low–abundance ions to an equal level. The resulting three–dimensional matrix (peak indices, samples and variables) was further subjected to multivariate data analysis. Partial least squares discriminant analysis (PLS–DA) was carried out to investigate and visualize the pattern of metabolite changes. This analysis was applied to obtain an overview of the complete dataset and discriminate those variables that are responsible for variation between groups. The PLS–DA model was evaluated through a cross–validation (R^2^ and Q^2^ parameters). The differential features were selected according to the VIP (variable importance in projection) list obtained from PLS–DA analysis. The PLS–DA was combined with univariate analysis (one–way ANOVA) with a P–value ≤ 0.05 to find differentially expressed metabolites. Using the Volcano Plot (VP) approach, which measure differentially accumulated metabolites based on *t* statistics and fold changes simultaneously, we also highlighted the metabolites with a −1 ≤ logFC ≥ 1 and statistically significant difference (P–value < 0.05) between genotypes at each time point. Identification of putative metabolites was performed using accurate mass metabolites reported in different publicly available databases such as METLIN, KEGG, Pubchem, HMDB and Plant Metabolic Network. Additionally, further partial identification of the most significant metabolites was made by comparison of MS/MS fragmentation patterns against reference compounds found in databases such as METLIN.

## Results

### Validation of Met.II.15 QTL and high–resolution mapping using HIFs

We initially focused on the GSL related phenotype of the Met.II.15 QTL to begin identifying the causal gene. However, the Met.II.15 QTL does not overlap any known biosynthetic or regulatory GSL genes. Thus, to facilitate map–based cloning of Met.II.15, we first validated it using HIF focusing on GSL phenotype (Fig.1A). To avoid temporal metabolite variation, in all the phenotype experiments, samples for GSLs analysis were harvested at mid-day. An initial screen of 200 plants from the HIF364 [Bay-0 × Sha] progeny, comparing plants homozygous for the Sha allele with plants homozygous for the Bay-0 allele and heterozygous at the QTL region, confirmed the phenotypic impact of Met.II.15 QTL on GSLs accumulation (Fig. 1B; Table S2). As previously found in the QTL mapping, total and individual aliphatic and indolic GSLs content of Met-HIF364 [Sha] lines was consistently higher than those of Met–HIF364 [Bay] at marker MSAT2.28 where the QTL was mapped originally. Lines heterozygous for HIF364 showed intermediate aliphatic and total GSL phenotypes, suggesting a semi–dominant relationship between the Bay-0 and Sha alleles (Fig. 1B). We then screened 1600 F2 plants and identified 32 recombination events in the initial heterozygous region of ~6 Mb between markers MSAT2008, MSAT2.28 and IND628. By recurrent genotypic selection and phenotypic analysis of recombinants within the QTL interval we selected eight lines since they all had a recombination break point between MSAT2.28 and IND628 and segregated for the GSL genotype (Fig. 1C; Table S3). Fine–mapping of Met.II.15 by adding new markers in the target region and progeny testing of lines homozygous for the different recombinant alleles, HIF364 [Bay x Sha] F3, narrowed the Met.II.15 locus to the region between markers M.II.D and M.II.E (Fig. 1C, D). Five recombinant lines (R4–R8) and control parental lines homozygous for Sha in the target region showed increased GSL content. In contrast, three recombinant line (R1–R3) and control parental line homozygous for Bay in the target region showed decreased GSL content (Fig. 1D).

**Figure 1.**
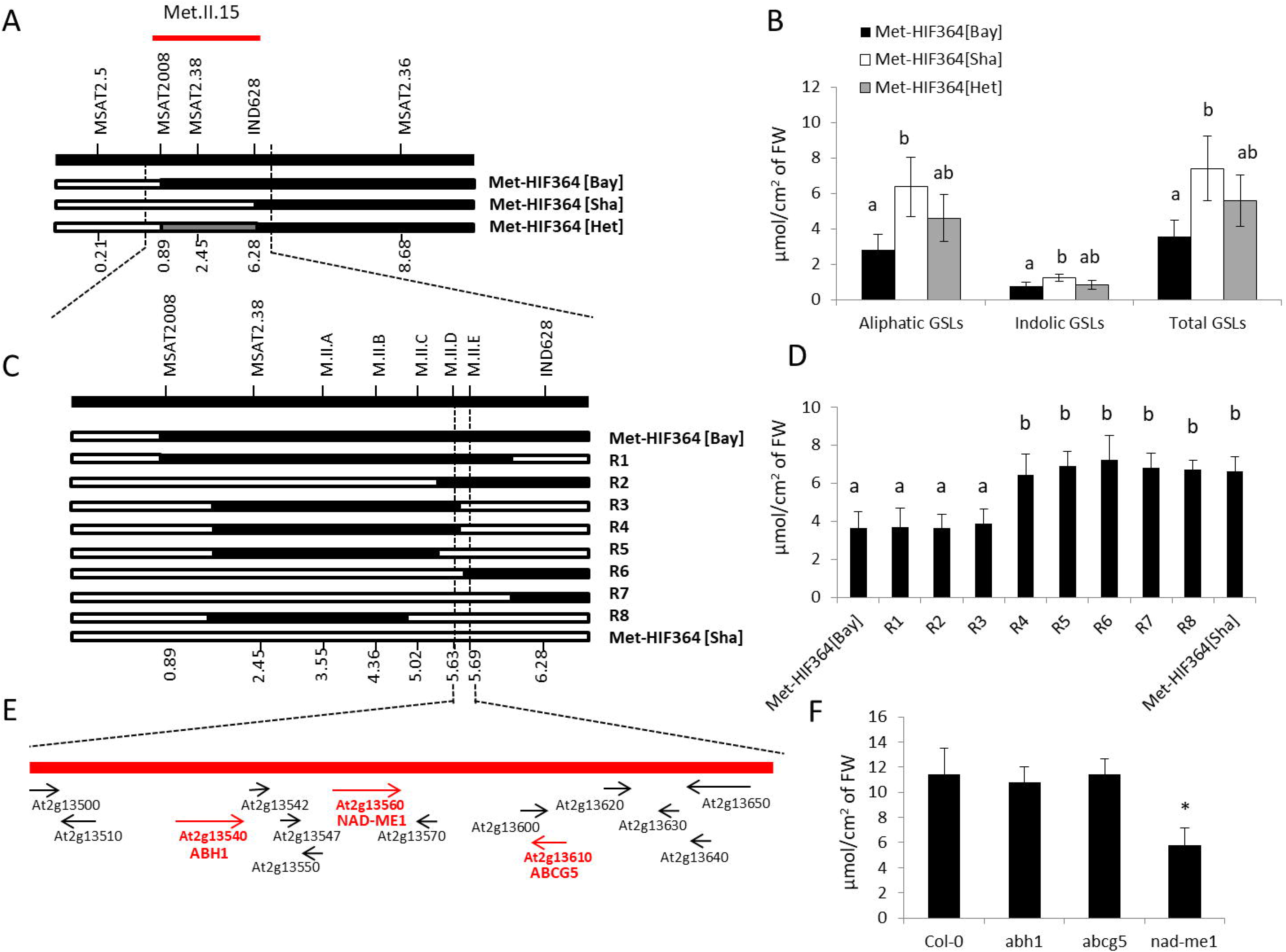
Fine mapping, phenotype characterization of heterogeneous inbred families (HIFs) segregating for Met.II.15 and candidate gene analysis. **A)** Map location of Met.II.15 QTL in a region (at MSAT2.38 and adjacent markers) on Arabidopsis chromosome 2 based on Bay x Sha QTL analysis (*25–27*). Molecular markers are represented with their corresponding names and positions in megabases below according to TAIR (see Supplemental Table1). **B)** Glucosinolate (GSL) (μmol/cm^2^ of FW) phenotypes from F2 plants (n=200) from HIF364 [Bay x Sha] (See table S2). **C)** High-resolution linkage map of selected F2 recombinant HIF364 (R1-8) narrowed the Met.II.15 locus to the region between markers M.II.D and M.II.E. **D)** Total GSLs progeny testing of fixed recombinant plants (HIF364 [Bay x Sha] F3, n=24). Five recombinant lines (R4-R8) and control parental lines homozygous for Sha in the target region showed increased GSL content. In contrast, three recombinant line (R1-R3) and control parental line homozygous for Bay in the target region showed decreased GSL content (See Table S3). **E)** Predicted protein coding genes in the region based on Arabidopsis reference genome (TAIR). Candidate genes with variation on transcript abundance between Bay-0 and Sha alleles are highlighted in red. **F)** Average GSL accumulation (μmol/cm^2^ of FW) of the evaluated T-DNA insertion lines from candidate genes (n=24) (See Table S4). Different letters indicate statistically significant differences determined by analysis of variance (ANOVA) with post□Jhoc Tukey’s honestly significant difference (HSD) test (P < 0.05). Error lines represent ± standard deviation of the mean.

### Identification of a candidate gene for the Met.II.15 QTL

The chromosomal region delimited by the M.II.D and M.II.E interval in the Met.II.15 region confidence interval includes 14 predicted protein coding genes (Fig. 2E; Table 1). Accordingly to previous Bay-0 × Sha eQTL analysis (*26*) we anticipate natural variation in gene expression for Met.II.15 causative gene. Thus, we focused only on the genes with evidence for a cis–eQTL. This analysis reduced the candidates to three genes: *ABA HYPERSENSITIVE 1* (At2g13540; *ABH1*), *NAD-DEPENDENT MALIC ENZYME 1* (At2g13560; *NAD–ME1*) and *ABC TRANSPORTER G FAMILY MEMBER 5* (AT2g13610; *ABCG5*). From them, in agreement with the overlapping circadian network, *NAD–ME1* and *ABCG5* had also a diel expression pattern (*40*). To confirm the phenotypic effects, GSL levels were evaluated in T–DNA insertion lines from each of the candidate genes with cis–eQTL (Fig. 1F; Table S4). Results evidenced that only *NAD–ME1* knockout mutant showed significant effects on GSLs levels. Simultaneously, we constructed co–expression networks by searching for connections between candidates and genes that are affiliated with GSL metabolism and circadian clock. Results corroborated that only *NAD–ME1* knockout mutant tightly co–expressed with SUR1 and SUR2, central components of the GSLs biosynthetic pathway. Moreover, *NAD–ME1* co–expressed with CRY2 (Cryptochrome 2) and PHYA (Phytochrome A), key components of the circadian oscillator complex (*41*). Thus, we hypothesize that *NAD–ME1* is the most likely candidate gene underlying the Met.II.15 QTL.

**Figure 2.**
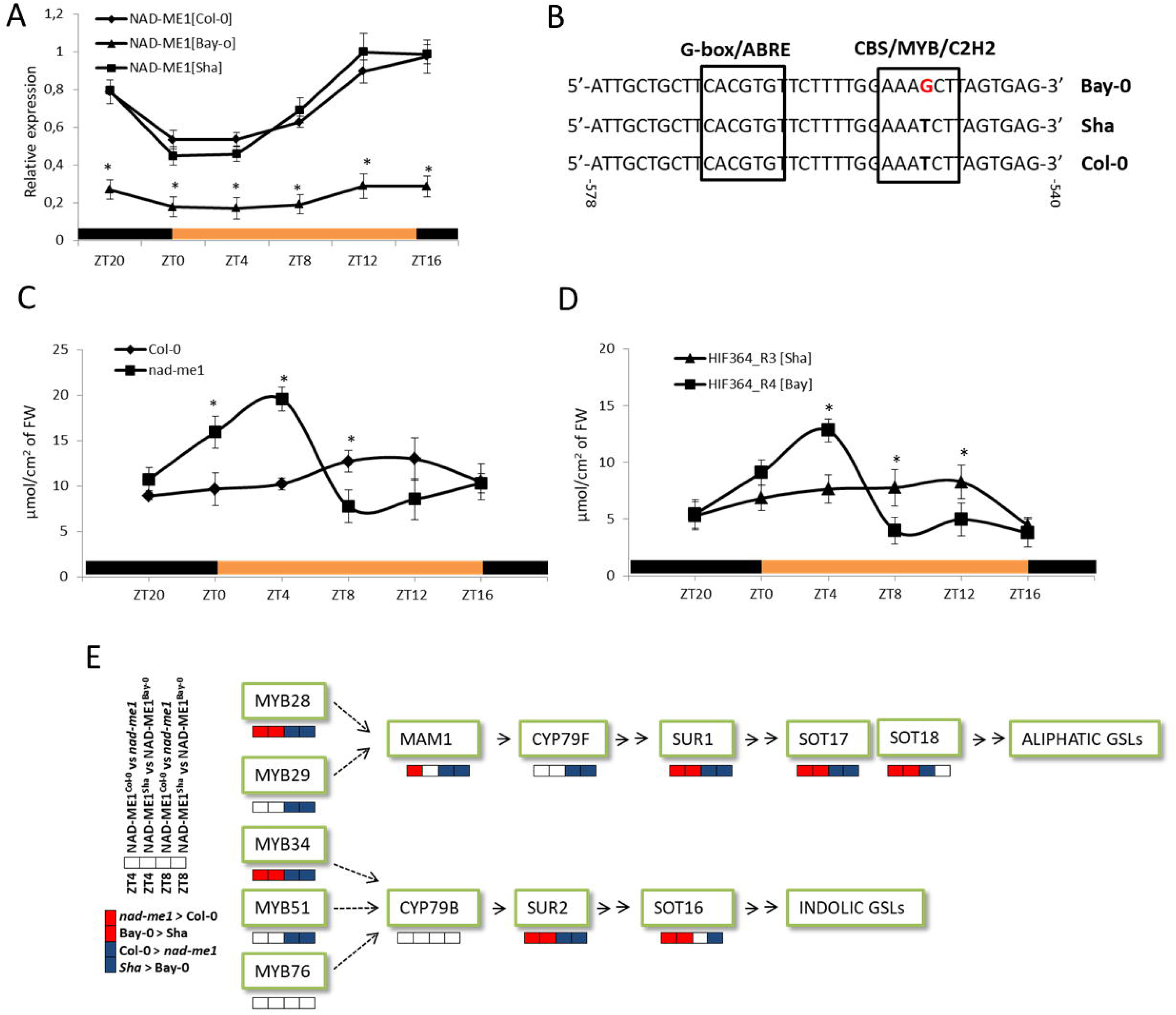
NAD-ME1 allelic effect and promoter analysis. **A)** Diurnal expression pattern of NAD-ME1 mRNA in long day (LD) conditions in three-weeks-old Arabidopsis leaves from three different genotypes: HIF364_R3 (NAD-ME1^Bay-0^), HIF364_R4 (NAD-ME1^Sha^) and Col-0 (NAD-ME1^Col-0^). Samples were harvested at various Zeitgeber times (ZT) every 4h in a 24h time-course experiment. **B)** Predicted cis-acting regulatory elements by PLACE in the NAD-ME1 promoter region are shown in black box. Sequence variation between Bay-0, Sha and Col-0 genotypes is highlighted in red. Numbers indicate positions from putative transcription start site. Sequence between −540 and −578 is shown, with two identified transcription factor binding motifs, one G-box/ABRE and one CBS/MYB/C2H2 binding site. **C, D)** Diurnal total glucosinolate content (μmol/cm^2^ of FW) of three-weeks-old Arabidopsis leaves from four different genotypes: NAD-ME1^Col-0^ vs *nad-me1* knockout mutant and NAD-ME1^Bay-0^ vs NAD-ME1^Sha^. Samples were harvested at various ZT every 4h in a 24h time-course experiment. **E)** Expression data of GSL aliphatic and indolic biosynthetic pathway genes from leaves of NAD-ME1^Col-0^ vs *nad-me1* knockout mutant and NAD-ME1^Bay-0^ vs NAD-ME1^Sha^ genotypes at ZT4 and ZT8. Highlighted with red squares are significantly up-regulated genes from NAD-ME1^Bay-0^ and *nad-me1* knockout mutant; Highlighted with blue squares are significantly up-regulated genes from NAD-ME1^Sha^ and NAD-ME1^Col-0.^ In all experiments three individual plants per genotype from two independent experiments were harvested each time point. Error lines represent ± standard deviation of the mean. The asterisk indicates statistically significant differences between mean values according to Student's t test (*p < 0.05).

**Table 1.**
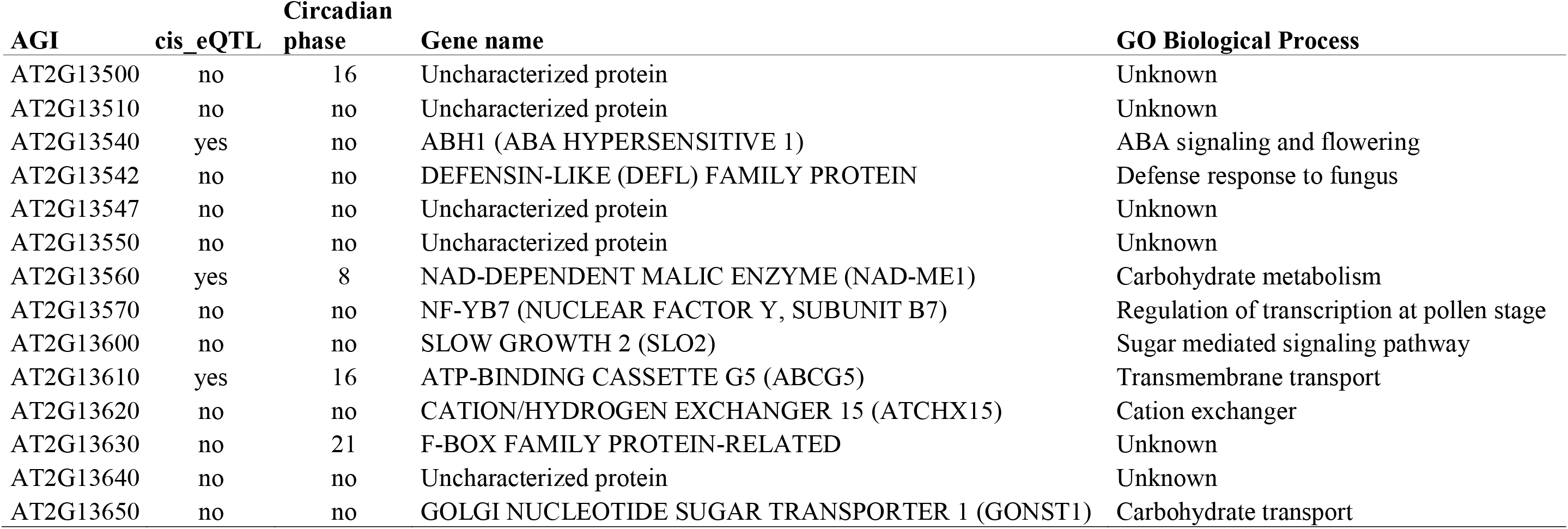
Shown is the list of candidate genes in the interval between markers M.II.D and M.II.E for the Met.II.15 QTL (At2g13500 to At2g13650) that have either a cis-eQTL or are regulated in a circadian manner with the phase reported.

| AGI | cis_eQTL | Circadian<br>phase | Gene name | GO Biological Process |
| --- | --- | --- | --- | --- |
| AT2G13500 | no | 16 | Uncharacterized protein | Unknown |
| AT2G13510 | no | no | Uncharacterized protein | Unknown |
| AT2G13540 | yes | no | ABH1 (ABA HYPERSENSITIVE 1) | ABA signaling and flowering |
| AT2G13542 | no | no | DEFENSIN-LIKE (DEFL) FAMILY PROTEIN | Defense response to fungus |
| AT2G13547 | no | no | Uncharacterized protein | Unknown |
| AT2G13550 | no | no | Uncharacterized protein | Unknown |
| AT2G13560 | yes | 8 | NAD-DEPENDENT MALIC ENZYME (NAD-ME1) | Carbohydrate metabolism |
| AT2G13570 | no | no | NF-YB7 (NUCLEAR FACTOR Y, SUBUNIT B7) | Regulation of transcription at pollen stage |
| AT2G13600 | no | no | SLOW GROWTH 2 (SLO2) | Sugar mediated signaling pathway |
| AT2G13610 | yes | 16 | ATP-BINDING CASSETTE G5 (ABCG5) | Transmembrane transport |
| AT2G13620 | no | no | CATION/HYDROGEN EXCHANGER 15 (ATCHX15) | Cation exchanger |
| AT2G13630 | no | 21 | F-BOX FAMILY PROTEIN-RELATED | Unknown |
| AT2G13640 | no | no | Uncharacterized protein | Unknown |
| AT2G13650 | no | no | GOLGI NUCLEOTIDE SUGAR TRANSPORTER 1 (GONST1) | Carbohydrate transport |

Our candidate, *NAD–ME1* gene, encodes a malic enzyme (ME) (EC 1.1.1.39) found in mitochondria of all cells (*42*) . MEs are widespread in all kinds of organisms and catalyzes the reversible oxidative decarboxylation of malate to pyruvate, CO2, and NAD(P)H in the presence of a divalent metal ion (*43*). It has been reported that MEs play a central role in the management of flux through the TCA cycle (*44*). However, to our knowledge *NAD–ME1* gene was not related before with GSL metabolism and/or circadian rhythmicity. Hence, in the next sections, several experiments were carried out in order to elucidate the interplay between plant metabolome and daily rhythms mediated by *NAD–ME1*.

### *NAD–ME1* alleles show altered diel regulation

To measure the potential influence of diurnal variation in conjunction with genetic variation, we studied the oscillation patterns of *NAD–ME1* gene expression in the progeny of the recombinant lines R3 (*NAD–ME1*^Bay-0^) and R4 (*NAD–ME1*^Sha^) carrying either parental allele in the final target region as well as in the Col-0 natural accession (*NAD–ME1*^Col-0^). RT*–*qPCR showed that the *NAD–ME1*^Sha^ and *NAD–ME1*^Col-0^ transcript levels are differently expressed between day and night (Fig. 2A). *NAD–ME1* transcript levels from Col-0 and Sha alleles exhibited their peaks from evening to early night and troughs at dusk suggesting circadian clock regulation. In contrast *NAD–ME1*^Bay-0^ transcripts were significantly lower than those of *NAD–ME1*^Sha^ and *NAD–ME1*^Col-0^ and lost the time*–*of*–*day*–*specific peak expression.

Further sequencing of the Bay-0 and Sha promoter and body gene of *NAD–ME1* revealed 51 SNPs and one deletion between Bay and Sha allele (Fig. S2). There were three SNPs within the predicted body of the RNA but these did not affect any key splicing sites or amino acid change. Thus, it is more likely that cis*–*eQTL is caused by the variation between the Bay and Sha promoters. To investigate that, *NAD–ME1* promoter sequences were analyzed through online program PLACE (www.dna.affrc.go.jp/PLACE/). Results showed that the 1.6 kb sequence upstream of the translation initiation codon ATG contains 20 types of putative cis-acting elements (Table 2). Supporting the idea that the clock regulates *NAD–ME1* transcripts, we found cis-regulatory elements known as the evening element (EE; AAAATATCT), which was identified in the promoters of Arabidopsis clock*–*controlled genes that are co*–*expressed late in the day (*9*), one putative circadian regulation element (CAANNNNATC) and two *CIRCADIAN CLOCK ASSOCIATED 1* (*CCA–1)* binding sites (CBS; AAATCT). Interestingly, we found that within 546 bp of the transcription start site a SNP in *NAD–ME1* Bay-0 allele promoter sequence may disrupt the putative CBS/MYB motif (Fig. 2B). It is remarkable that in close proximity to this sequence, is localized the G*–*box core element (Fig. 2B; Table 2). Typically, G*–*box and CBS motifs are part of light/circadian regulated gene promoters and usually cooperate in defining the transcript oscillation properties (*45*). In addition, disrupted sequence in Bay allele may be also affecting the binding of C2H2 zinc-finger proteins which are involved in participate in various aspects of normal plant growth and development, as well as in environmental stress tolerance regulation (*46*). Although further work is required to identify the specific transcription factor that leads to the alterations in *NAD–ME1* gene expression, we hypothesized that difference of *NAD–ME1* expression among alleles is likely to be caused by polymorphisms in cis*–*acting elements at the promoter.

**Table 2.**
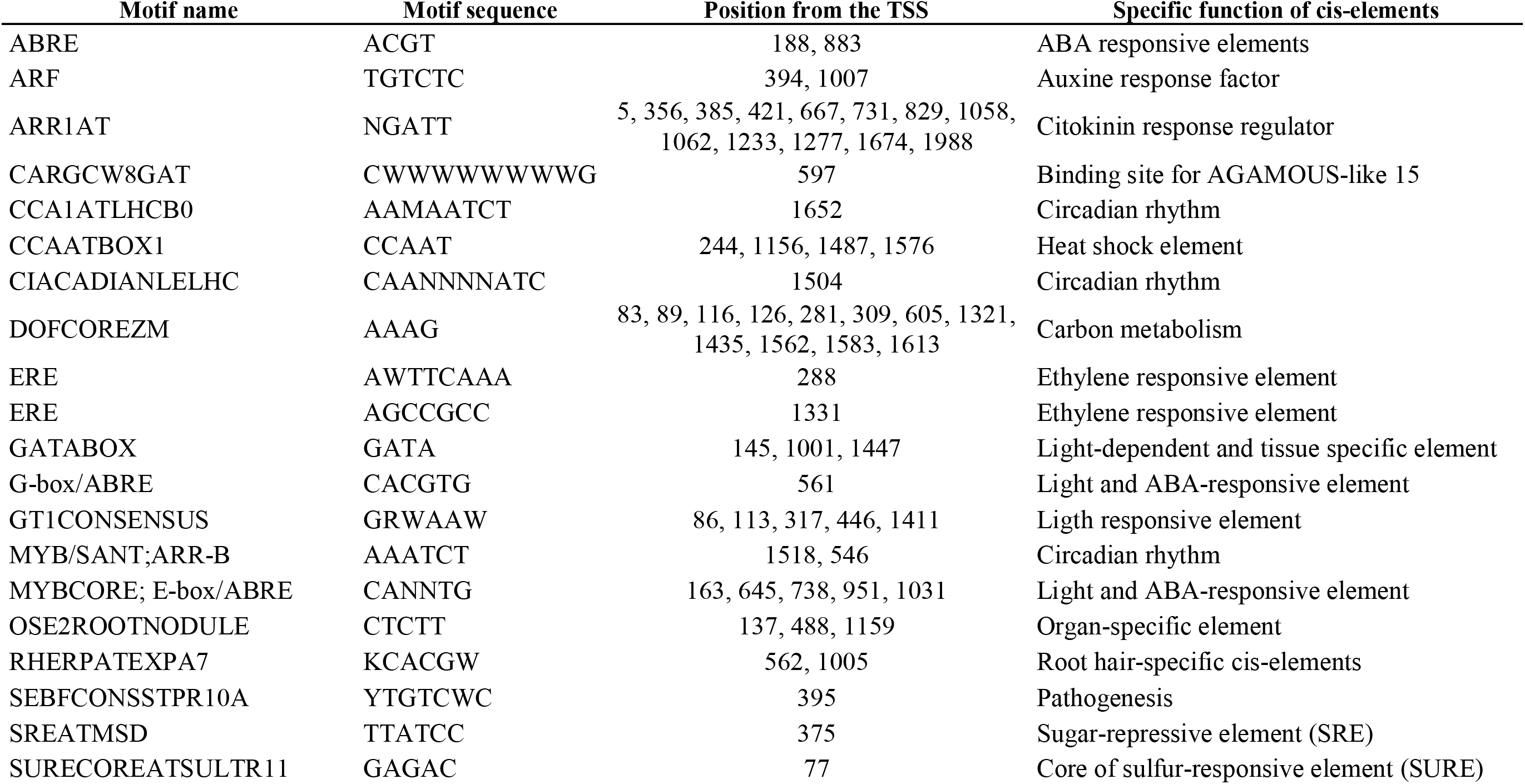
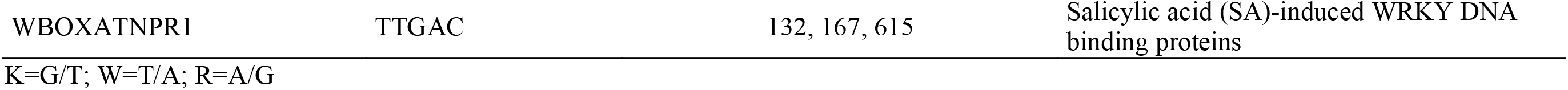
List of known cis-element motifs found in an interval of 1.6 kb upstream from the transcription start site (TSS) of NAD-ME1 promoter in the Bay-0 and Sha alleles. Motifs are listed alphabetically including their motif name, sequence, position from the TSS and specific function.

### Amplitude of the GSLs diel variation was affected by *NAD–ME1* alleles

The finding that *NAD–ME1* transcripts could be regulated in a diel manner prompted us to test if this also leads to an alteration in the temporal variation in GSL accumulation during the day. To test if *NAD–ME1* controls diel variation in GSL metabolism, we carried out a time course under LD cycle conditions in three*–*weeks*–*old plants, sampling every 4 hours over 1 day in the *NAD–ME1*^Bay-0^ and *NAD–ME1*^Sha^ genotypes, as well as in the knockout mutant line of the candidate gene *NAD–ME1* (*nad–me1*) and in the wild type, *NAD–ME1*^Col-0^. Results showed that the amplitude of the GSLs diel variation was higher in the *nad–me1* and *NAD–ME1*^Bay-0^ genotypes compared with *NAD–ME1*^Col-0^ and *NAD–ME1*^Sha^ alleles respectively (Fig. 2C, D). In general, during the first hours of light, with a maximum at ZT4, the *nad–me1* mutants and *NAD–ME1*^Bay-0^ accumulated significantly more GSLs than *NAD–ME1*^Col-0^ and *NAD–ME1*^Sha^ lines respectively. On the contrary, at mid-day (ZT8) GSL content in *nad–me1* mutant and *NAD–ME1*^Bay-0^ lines showed a clear reduction in GSL content. Therefore, different alleles of *NAD–ME1* affect the amplitude of GSLs diel variation.

To further investigate the influence of *NAD–ME1* on GSLs accumulation, we studied transcript levels of GSL related genes at ZT4 and ZT8 (Fig. 2E; Table S5). Regulation of numerous GSL biosynthesis genes was strongly affected in the *nad–me1* mutant and in the recombinant line caring Bay allele. In general, down-regulation of *NAD–ME1* transcript levels was related with up*–*regulation of GSL related transcript genes during morning while dampening during the evening. Thus, expression of *NAD–ME1* is correlated with the GSL metabolism suggesting that *NAD–ME1* is a strong candidate for Met.II.15 cis*–*eQTL.

### Untargeted metabolomics revealed that *NAD–ME1* loss of function induces time–specific metabolome changes in Arabidopsis

Since *NAD–ME1* was assumed to play a central role in the management of flux through the TCA cycle by coordination of the carbon and nitrogen metabolisms in Arabidopsis (*47*) and Met.II.15 co-localizes with a major QTL controlling central metabolism associated with the TCA cycle (*25*), we suggest that *NAD–ME1* may be also affecting broad metabolomics changes. Thus, we carried out an untargeted metabolomics analysis using UPLC*–*Q*–*TOF*–*MS/MS to evaluate the effect of *NAD–ME1* loss of function on plant metabolome. We analyzed global trends of metabolite variations by searching for changes that occur in leaves of *nad–me1* mutants compared with Col-0 at ZT4 and ZT8. These time points were chosen because wider amplitude on GSLs variation was found in previous sections. PLS-DA score plots showed a clear separation between Col-0 and *nad–me1* group plant genotypes (Fig. 3A). We detected 57 and 81 differentially expressed metabolites among *nad–me1* mutant and Col-0 at ZT4 and ZT8, respectively (Table S6). Among the 138 ions significantly altered, only 15 were coincident at both time points studied (Fig. 3B). Thus, close to 90% of the significant altered ions among genotypes, including TCA intermediates, were dependent upon the day*–*time, which evidenced complex temporal metabolomics changes in Arabidopsis *NAD–ME1* network (Fig. 3C). Metabolomics comparison of *NAD–ME1*^Bay-0^ and *NAD–ME1*^Sha^ genotypes corroborated the temporal shift on primary metabolites related with TCA cycle (Fig. S3). At ZT4, the time point equivalent to that used for the QTL mapping sample, the direction of effect agreed with the previously published analysis from Met.II.15 in the Bay × Sha RIL population (*25*). This suggests that *NAD–ME1* plays a role in mediating the temporal partitioning of TCA intermediates within the Arabidopsis leaf.

**Figure 3.**
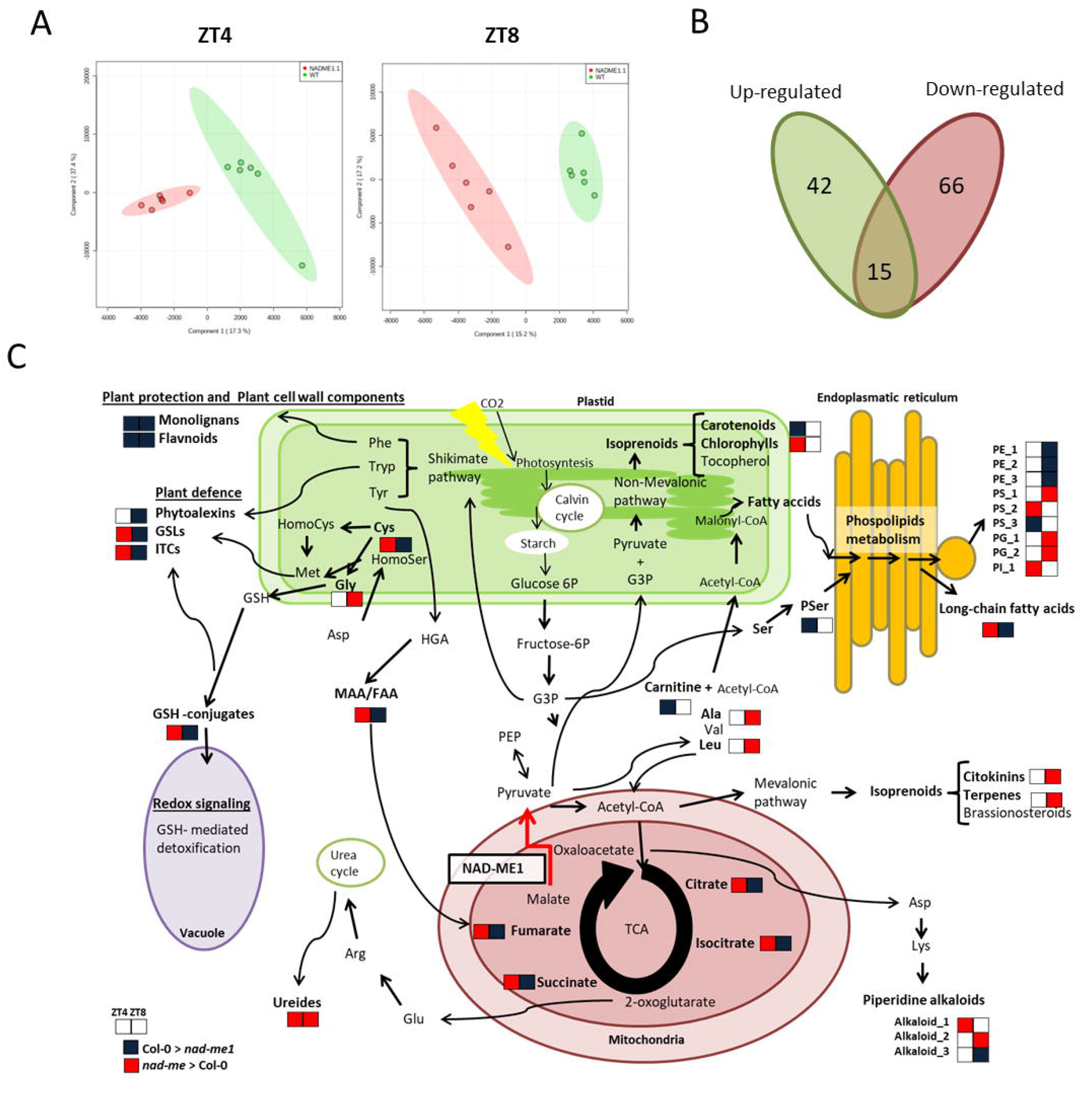
Impact of NAD-ME1 in Arabidopsis metabolome. **A)** PLS-DA score plots from UPLC-Q-TOF-MS/MS analysis showing a clear group separation between Col-0 and *nad-me1* at two different time points (ZT4 and ZT8). Green and red ellipsoids show the 95% confidence interval in Col-0 and *nad-me1* plants, respectively. **B)** Comparison of the significant altered ions at both time points analyzed. **C)** Schematic model of NAD-ME1 metabolomics changes in the cell. Red arrow represents the enzymatic step catalyzed by NAD-ME1. Highlighted with red squares are significantly up-regulated metabolites from *nad-me1* knockout mutant; Highlighted with blue squares are significantly up-regulated metabolites from Col-0. G3P, 3-phosphoglycerate; PEP, phospho*enol*pyruvate; Fructose-6P, fructose 6-phosphate; Glucose 6-P, glucose 6-phosphate; IAA, indole-3-acetic acid; JA, jasmonic acid; GSLs, glucosinolates; ITCs, isothiocyanates; GSH, glutation; Phe, phenlylalanine; Tyr, tyrosine; Tryp, tryptophan; Cys, cystenine; Met, methionine; Gly, glycine; Asp, asparagine; Ser, serine; PSer, Phosphoserine; Val, valine; Leu, leucine; Arg, arginine; Glu, glutamic acid; Lys, lysine; PE, Phosphatidylethanolamine derivative; PS, Phosphoserine derivative; PG, Phosphatidylglycerophosphate derivative; PI: Phospho-myo-inositol derivative. For complete metabolite information see Table S6.

Rather than TCA intermediates itself, lack of *NAD–ME1* caused alterations in many other metabolites involved in a wide a wide range of metabolic processes (Table S6). At ZT4, in terms of fold change, the major differences among genotypes were found for isoprenoids potentially involved in plant photosynthesis and cell-wall precursors. At ZT8, amino acids and their derivatives were the most altered compounds. In addition, several compounds related with phospholipid metabolism, fatty acids, cysteine and glutathione conjugates, flavonoids, and plant immunity compounds (phytoalexins, isothiocyanates and alkaloids) were altered in the mutant in a time*–*dependent fashion (Fig. 3C; Table S6).

### *NAD–ME1* regulatory network construction

To understand the molecular mechanisms behind the *NAD–ME1* effect on plant metabolism, we worked to identify potential co*–*regulated modules of our candidate gene which may explain metabolic changes between genotypes. To achieve that, we first searched for genes closely expressed with *NAD–ME1* through the ATTEDII co-expression database and then filtered for genes with a cis–eQTL in the Bay-0 × Sha RIL population (*26*). Selected genes were used as bait genes to construct the co*–*expression network (Fig. 4; Table S7).

**Figure 4.**
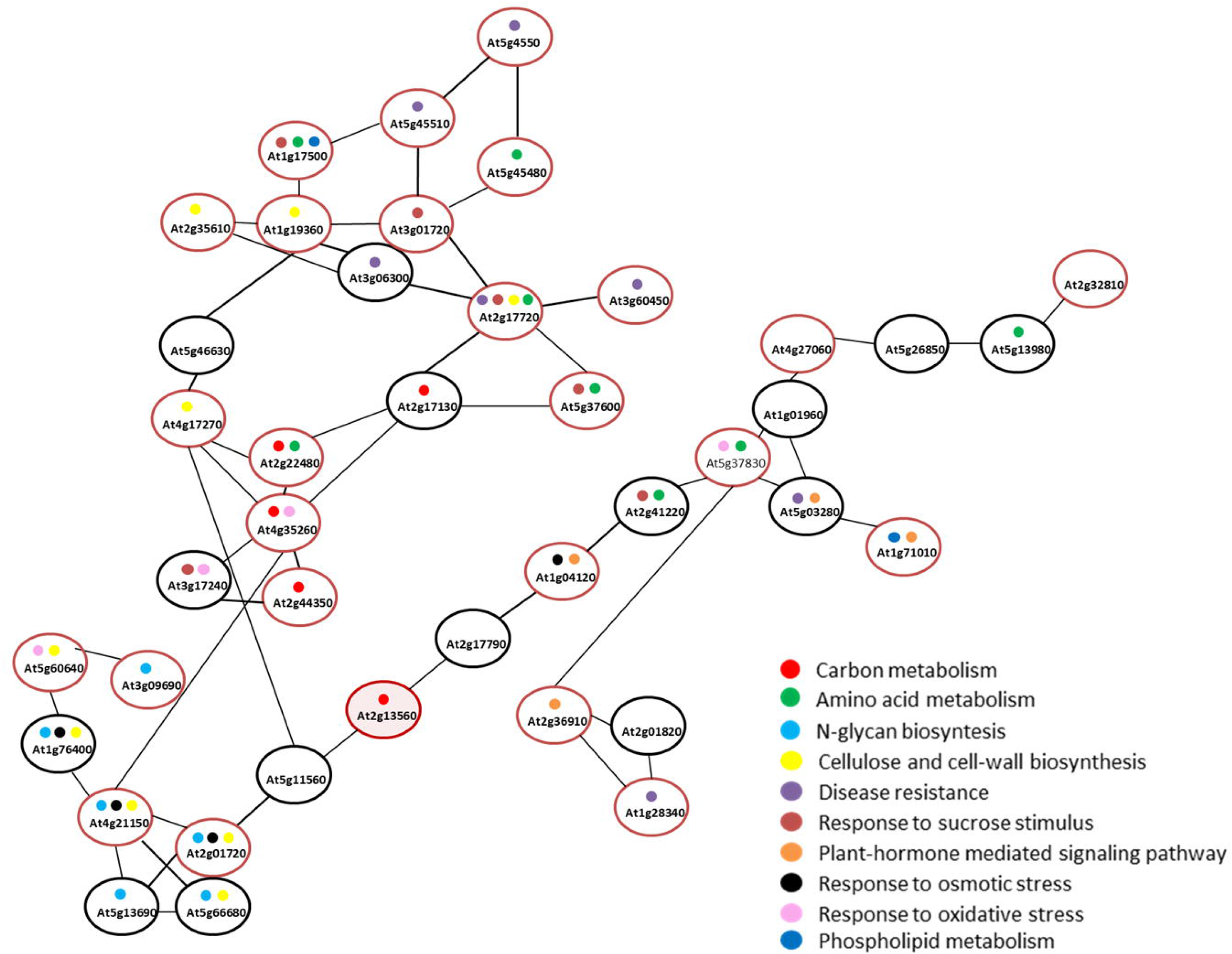
Co-expressed NAD-ME1 network. To construct the network, we filtered the first 30 genes correlated with *NAD–ME1* from the ATTED-II database. Among them, we selected only the genes with a cis–eQTL in the Bay-0 × Sha RIL population (*26*). The selected genes were used as bait genes. Then, co–expression relationship between the bait genes and their directly–connected genes were used to construct a network visualized using NetworkDrawer. Genes were subsequently interpreted by functional enrichment analysis based on Gene Ontology (GO) to identify and rank overrepresented functional categories in the network. Red circles depict genes with cis–eQTL. For complete gene information see Table S7.

Studying Gene Ontology (GO) terms in the ‘biological process’ annotation, revealed over*–*representation of genes involved in primary carbon metabolism, response to stimulus, amino acid metabolism, N*–*glycan biosynthesis, cell*–*wall remodeling and phospholipid metabolism . Remarkably, 73% of the genes from the *NAD–ME1* co-expressed network with an eQTL were assigned to plant response to different stimulus. This set included several genes shown to be involved in cell redox homeostasis, plant hormone signal transduction, response to sucrose stimulus and systemic acquired resistance related genes (Table S7). The largest hubs within the network were the *PROLYL 4-HYDROXYLASE 5*(*P4H5*; At2g17720), the *ISOCITRATE DEHYDROGENASE 1* (*IDH1*; AT4g35260) and *HAP6* (At4g21150). The *P4H5* catalyze an important post-translational modification required for proper cell-wall self-assembly and protein stability of transcription factors in response to biotic and abiotic plant stresses and hence involved in plant defense (*48*). The *IDH1* performs an essential role in cellular defense against oxidative stress-induced damage of the TCA cycle (*49*). Lastly, the *HAP6* gene is a member of the oligosaccharyltransferase (OST) multi*–* membrane protein complex, with a role in protein N*–*linked glycosylation and involved in cellulose biosynthesis in response to abiotic stress (*50*). All together, these results suggest that impaired *NAD–ME1* expression is correlated with various plant cell stress-sensing mechanisms affecting a combination of different signal transduction pathways.

## Discussion

During the last decade, increasing evidence suggest the existence of a complex interplay between plant metabolic pathways and daily accumulation of a large range of plant metabolites, including GSLs (*16–20, 27, 32*). However, identification and cloning of new loci controlling quantitative traits related to both plant metabolism and diel patterns have been less studied. In this work, we created NILs developed from HIFs that only segregates at a small region around the Met.II.15 QTL controlling temporal variation in primary and specialized metabolism in the Arabidopsis Bay-0 × Sha population. This approach facilitated map-based cloning and successfully identified *NAD–ME1*, involved in the decarboxylation of malate to pyruvate, as the most likely candidate gene underlying the Met.II.15 QTL leading to temporal shifts of TCA intermediates, GSL accumulation and altered regulation of several GSL biosynthesis pathway genes in a time-dependent manner.

The availability of specific recombinants of RILs containing *NAD–ME1* allelic variation allowed us to confirm the light/circadian dependent cis*–*eQTL on transcript abundance (Fig.2A). Transcripts of *NAD–ME1* from Col-0 and Sha alleles peaks from evening to early night, corroborating a major role of *NAD–ME1* during night mitochondrial metabolism (*51*). Transcripts from Bay-0 allele were significantly down*–*regulated and lost the time*–*of*–*day*–*specific peak expression, indicating that these differences in expression among alleles are likely to be caused by polymorphisms in light/clock*–* controlled cis*–*acting elements at the promoter. Interestingly, a polymorphism disrupting a putative CBS motif in *NAD–ME1* promoter region of Bay allele, may be potentially affecting the binding of the core clock elements *CCA1* and *LHY*. Supporting our findings, Graf et al. (*12*) found that *NAD–ME1* transcripts and protein levels are significantly reduced in *cca1/lhy* double mutants. These evidences suggested that *CCA1* and *LHY* are highly likely to regulate *NAD–ME1* expression.

Intriguingly, the loss of *NAD–ME1* transcript regulation was linked to temporal shifts of several TCA intermediates (Fig. 3; Fig S3; Table S6). Although the biological meaning of the linkage between circadian rhythms and TCA cycle remains unclear, Arabidopsis clock mutants showed deregulation of TCA metabolic homeostasis within plant mitochondria (*11, 52*). More recently, it has been reported that many TCA intermediates drove the diel changes under different natural and artificial light regimes in Arabidopsis (*15*). In general, these metabolites are low at dawn and high towards the end of the light period. However, we found that plants lacking *NAD–ME1* displayed enhanced levels of several TCA cycle intermediates after dawn and decreased at evening (Fig,3C; Fig. S3; Table S6). Interestingly, neither malate nor pyruvate, the substrate and product of the NAD–ME1 reaction, were altered in plants lacking *NAD–ME1* at the time points tested. This can be explained because *NAD–ME1* is not the sole source of pyruvate in mitochondria. In accordance with this, the other NAD-dependent malic enzyme *NAD-ME2*(At4g00570) and two genes encoding *MITOCHONDRIAL MALATE DEHYDROGENASE* (*MMDH*; At1g53240 and At3g47520) can metabolize malate in the mitochondria. It has been suggested that MMDHs and NAD-MEs manage the flux of malate through the TCA cycle differentially during a diurnal cycle. MMDHs would have a prevalent role during the light period, while NAD-MEs would be more important during the night period (*51*). This is consistent with the suggested role of *NAD–ME1* during night mitochondrial metabolism based on our results in time*–*of*–*day*–*specific peak expression (Fig. 2). When *NAD–ME1* is not present, like in *nad–me1* mutant plants used in the present study, Arabidopsis leaves display half of total NAD-ME enzyme activity compared with wild*–*type, presumably associated with NAD-ME2 activity (*51*). Therefore, we speculated that reduced NAD-ME activity during the dark period in Bay-0 and *nad–me1* produces an excess of mitochondrial TCA intermediates at the end of the night and, in consequence, many primary and secondary plant metabolites derived from TCA will be affected in a time*–*dependent fashion. Thus, *NAD–ME1* seems to be key factor involved in to an intricate system to modulate the flux into and out of central metabolism.

TCA cycle intermediates provide the carbon skeletons to support the biosynthesis of the majority of amino acids (*53, 54*). In the current work, we found that amino acids and related metabolites were altered in plants lacking *NAD–ME1*. Indeed, mutant plants displayed temporal shifts of several plant defensive compounds derived from amino acids including GSLs, flavonoids, phytoalexins and alkaloids (Fig. 3, Table S6). To our knowledge, *NAD–ME1* has not previously been implicated in the production of plant defensive compounds. Thus, potential explanations for temporal shift of these compounds may be related with altered cysteine and glutathione conjugates pool sizes during the light hours due to the lack of *NAD–ME1* during nocturnal metabolism (Fig.3C, Table S6). Cysteine occupies a central position in plant metabolism because it is a reduced sulfur donor molecule involved in the synthesis of essential biomolecules such as the other proteinogenic amino acid methionine, vitamins, cofactors, and the antioxidant glutathione (Romero et al., 2014). Methionine is the amino acid precursor of aliphatic GSLs and glutathione plays a role as sulfur donor in GSLs formation. In addition, several intermediates of tryptophan derived metabolites including indole GSLs as well as intermediates from phytoalexins biosynthesis were altered in the *nad–me1* mutant. These findings are in line with our data from the co*–*expressed *NAD–ME1* network (Fig.4; Table S7). One of the most significant GO categories identified from the *NAD–ME1* network were genes involved in amino acid metabolism, including cysteine, glutathione and tryptophan related genes. Remarkably, transcript variation of most of those genes was related with plant response to plant hormones and plant*–* pathogen interactions with a role in the plant systemic acquired resistance response (Fig. 4, Table S7). This agrees with the fact that the regulation of several GSL biosynthesis pathway genes, including major transcriptional regulators of the aliphatic and indolic GSLs biosynthesis pathways in response to biotic and abiotic stress (*MYB28*, *MYB29*, *MYB34*) were strongly time*–*dependent affected in plants lacking *NAD–ME1*. These results may indicate that *NAD–ME1* down*–*regulation might be a decisive factor for basal and inducible levels of chemical defenses and thereby plant resistance. Further research is necessary to test whether temporal alterations in plant phytochemicals under biotic and abiotic stress might affect plant defense response in *nad-me1* mutant plants.

A closer examination of untargeted metabolomics analyses revealed that rather than TCA intermediates itself and plant defensive compounds, the lack of *NAD–ME1* caused alterations in many other metabolites involved in a wide a wide range of metabolic processes (Fig.3C; Table S6). Several precursors of cell*–*wall components and phospholipids, critical components of cell membranes, appear to be temporary altered in the mutant. Moreover, down*–*regulation of *NAD–ME1* produced reduction in photosynthetic pigments while chlorophyll degradation products increased during the first part of the day*–*light hours. Similar modifications on metabolite composition that occurred in *nad–me1* mutant plants were before associated with plant responses against high light, cold or nutrient stresses among others (*55–57*). In concordance with that, genes related with phospholipid biosynthetic process and cell*–*wall organization in response to cold, osmotic stress, cell redox homeostasis and sucrose stimulus were over*–*represented in the *NAD–ME1* co*–*regulated network (Fig.4; Table S7). Since no effect on growth or development are found for *NAD–ME1* mutant plants under the tested conditions (Fig. S1) it seems that *NAD–ME1* down*–*regulation may initiate several mechanisms of acclimation and adaptation establishing a new metabolic homeostasis that mimics plant responses to environmental stress conditions. Results from this work will facilitate future research investigating how *NAD–ME1* would respond to diverse and dynamic environmental cues.

As a conclusion, this work identifies *NAD–ME1* as a novel actor of plant metabolomics network and provides new insights into the modular behavior of biochemical processes and their diel regulation within Arabidopsis. Transcript variation of the *NAD–ME1* gene led to temporal shifts of TCA intermediates, GSLs accumulation and altered regulation of several GSL biosynthesis pathway genes. In addition, lack of *NAD–ME1* induced time*–*dependent changes of plant phytochemicals related to plant immunity, initiated a complicated re*–*organization of plant cell*–*wall components and it seems to be involved in photosynthesis optimization. These findings suggest that *NAD–ME1* may coordinate plant primary and secondary metabolism in a diel dependent manner. Because some metabolites modulate the plant clock, it is needed to elucidate the interaction mechanism between the biological clock system and metabolism such as in the TCA cycle or GSLs pathways. A key future step is to investigate whether diurnal regulation of plant metabolome mediated by *NAD–ME1* may affect plant physiology, adaptation and survival under environmental stress conditions.

## Acknowledgements

This effort was funded by the grant RTI2018-094650-J-I00 by the Spanish Ministry of Science, Innovation and Universities to Marta Francisco.

## Author contributions

M.F., D.K., and E.C. conceived and designed the experiments. M.F., V.M.R., P.S. and R.A. assisted with the plant trials setup; M.F., and R.A. conducted laboratory work; V.M.R. did the metabolomics analysis; M.F. analyzed the data and wrote the manuscript. All authors read and approved the final manuscript. The authors declare that they have no competing interests.

