## Supplementary material for "Fine–mapping identifies *NAD–ME1* as a candidate underlying a major locus controlling temporal variation in primary and specialized metabolism in Arabidopsis": Supplental figures

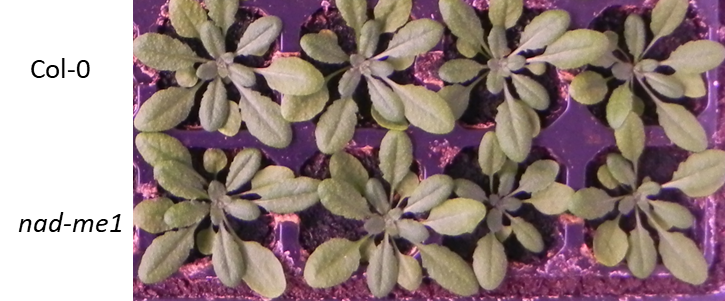

**Figure S1.** Arabidopsis plants from Col-0 and *nad-me1* genotypes after three weeks of germination grown under long day (LD) conditions.

```
Bay-0  1     TTCGAGAGTTCTTAGCTAAACAATCTGCTCGAGTATTAAAACTACGAGCTAATGCTGAAA  60
             ||||||||||||||||||||||||||||||||||||||||||||||||||||||||||||
Sha    1     TTCGAGAGTTCTTAGCTAAACAATCTGCTCGAGTATTAAAACTACGAGCTAATGCTGAAA  60

Bay-0  61    GAAGAGAAAAAACAAACTAAATACCAAAAAGAGGCCAATTCCGTTGCAAAAGTTGGTCAA  120
             ||||||||||||||||||||||||||||||||||||||||||||||||||||||||||||
Sha    61    GAAGAGAAAAAACAAACTAAATACCAAAAAGAGGCCAATTCCGTTGCAAAAGTTGGTCAA  120

Bay-0  121   TCGTCTTGGTGATCAACGAGAGAGATCAAATCTGCGCAATCGGTAGCAAAGGTATCTGTA  180
             ||||||||||||||||||||||||||||||||||||||||||||||||||||||||||||
Sha    121   TCGTCTTGGTGATCAACGAGAGAGATCAAATCTGCGCAATCGGTAGCAAAGGTATCTGTA  180

Bay-0  181   TCGCCAATGCTGATCATGCATTCCATTGCCCCTAGCAAAGATTCAAGTTCGACGTAAAGC  240
             ||||||||||||||||||||||||||||||||||||||||||||||||||||||||||||
Sha    181   TCGCCAATGCTGATCATGCATTCCATTGCCCCTAGCAAAGATTCAAGTTCGACGTAAAGC  240

Bay-0  241   GGAGATAAGTTTTTTCGAAGAAAAAAAAAGTTACTCATGTAAGCGAGTAAGTAACTCAAA  300
             ||||||||||||||||||||||||||||||||||||||||||||||||||||||||||||
Sha    241   GGAGATAAGTTTTTTCGAAGAAAAAAAAAGTTACTCATGTAAGCGAGTAAGTAACTCAAA  300

Bay-0  301   AAAACAAAAAAAAACCTAGTTTTCTCTCTCGCAAAACCAGCCGCCGTCTAAAGAGGTTTT  360
             ||||||||||||||||||||||||||||||||||||||||||||||||||||||||||||
Sha    301   AAAACAAAAAAAAACCTAGTTTTCTCTCTCGCAAAACCAGCCGCCGTCTAAAGAGGTTTT  360

Bay-0  361   GCAGTTTTGAGGCTAGAATGCGGTGGCTACCTTAGCACTGATTCTGGCCTTGTGCGTCTT  420
             ||||||||||||||||||||||||||||||||||||||||||||||||||||||||||||
Sha    361   GCAGTTTTGAGGCTAGAATGCGGTGGCTACCTTAGCACTGATTCTGGCCTTGTGCGTCTT  420

Bay-0  421   TGTTTAATTTCGTCTGTAGATTTCTTCCTTCTCGTTTTTATGGTGATATGGTTGTAGGGT  480
             ||||||||||||||||||||||||||||||||||||||||||||||||||||||||||||
Sha    421   TGTTTAATTTCGTCTGTAGATTTCTTCCTTCTCGTTTTTATGGTGATATGGTTGTAGGGT  480

Bay-0  481   TGATGGTGTTGGGGAGTGCGGGGACCAATTTCTCTTCTATATCTGTTTCGGAGGTTGGTC  540
             ||||||||||||||||||||||||||||||||||||||||||||||||||||||||||||
Sha    481   TGATGGTGTTGGGGAGTGCGGGGACCAATTTCTCTTCTATATCTGTTTCGGAGGTTGGTC  540

Bay-0  541   TCTATGTTGTTGGCAGCTCCCGTTTTTAGGCTTTGTTTCAATTTTGGCTGTATATGCGGC  600
             ||||||||||||||||||||||||||||||||||||||||||||||||||||||||||||
Sha    541   TCTATGTTGTTGGCAGCTCCCGTTTTTAGGCTTTGTTTCAATTTTGGCTGTATATGCGGC  600

Bay-0  601   TGGGGTTTTGATTGATTTCCAGATCTTGATCTGGTTTTGTCAGTTGAAGCTATGTCAGTG  660
             ||||||||||||||||||||||||||||||||||||||||||||||||||||||||||||
Sha    601   TGGGGTTTTGATTGATTTCCAGATCTTGATCTGGTTTTGTCAGTTGAAGCTATGTCAGTG  660

Bay-0  661   TTTGTCACGATATGGAATTGTTTCCTTTTGTCTTCCCCCGTTGTGTTCTGTGTGTATGCA  720
             ||||||||||||||||||||||||||||||||||||||||||||||||||||||| ||||
Sha    661   TTTGTCACGATATGGAATTGTTTCCTTTTGTCTTCCCCCGTTGTGTTCTGTGTGTTTGCA  720

Bay-0  721   GTTGAAGATGAAGTCGTTTTGAGTTGGTGGATGCCTCTGGTGATGGGTTCTGACCGGGTT  780
             |||||||||||||||||||||| |||||||||||||||||||||||||| ||||||||||
Sha    721   GTTGAAGATGAAGTCGTTTTGAATTGGTGGATGCCTCTGGTGATGGGTTTTGACCGGGTT  780

Bay-0  781   ATGTGACGTTCTTGTTGTGGACTAGGCGACGACTCGTCAGGGTGGTGGCTGTTTGGAGGC  840
             ||||||||||||||||||||||||||||||||||||||||||||||||||||||||||||
Sha    781   ATGTGACGTTCTTGTTGTGGACTAGGCGACGACTCGTCAGGGTGGTGGCTGTTTGGAGGC  840

Bay-0  841   GATTTAGGTTGTCTTCCGGTGCGGGTATTTGGCTGAAACTGGGTCGGCAAATGCATGCGT  900
             ||||||||||||||||||||||||||||||||||||||||||||||||||||||||||||
Sha    841   GATTTAGGTTGTCTTCCGGTGCGGGTATTTGGCTGAAACTGGGTCGGCAAATGCATGCGT  900

Bay-0  901   GCAAGAGGTGTTTGTCTTATTTGGCTTGTCATTTGGCGATTCTCAAGGGCTTCCCTTGTT  960
             ||||||||||||||||||||||||||||||||||||||||||||||||||||||||||||
Sha    901   GCAAGAGGTGTTTGTCTTATTTGGCTTGTCATTTGGCGATTCTCAAGGGCTTCCCTTGTT  960


Bay-0  961   CCTAGTTTATTTTCGTCTTCTCGTGTGCCTTGGTATGGTGTGATTGTTGTCTCTCTAGTA  1020
             |||||||||||||| |||||||||||||||||||||||||||||||||||||||||||||
Sha    961   CCTAGTTTATTTTCATCTTCTCGTGTGCCTTGGTATGGTGTGATTGTTGTCTCTCTAGTA  1020

Bay-0  1021  TCATTTGGTTCTTGCTTTCGTTCCTGGCGGTTTGACTTCAGAAAGATGTCTTAATTTTGT  1080
             ||||||||||||||||||||||||||||||||||||||||||||||||||||||||||||
Sha    1021  TCATTTGGTTCTTGCTTTCGTTCCTGGCGGTTTGACTTCAGAAAGATGTCTTAATTTTGT  1080

Bay-0  1081  GGCATTACCAGTCTTATTGCTGCTTCACGTGTTCTTTTGGAAAGCTTAGTGAGAACCTCT  1140
             ||||||||||||||||||||||||||||||||||||||||||| ||||||||||||||||
Sha    1081  GGCATTACCAGTCTTATTGCTGCTTCACGTGTTCTTTTGGAAATCTTAGTGAGAACCTCT  1140

Bay-0  1141  ATGGAACAATAGGTCAGCTTAGTGCTTTGGTCTCCATCTCTTCTTTTTGCGAGTTTATCT  1200
             |||||||||||||||||||||||||||||||||| |||||||||||||||||||||||||
Sha    1141  ATGGAACAATAGGTCAGCTTAGTGCTTTGGTCTCAATCTCTTCTTTTTGCGAGTTTATCT  1200

Bay-0  1201  TGTTGTTTTTACCTTTCTTGGTAATTTGGCTGTCTGGTTTTTAGGGATTAGTTTTCCTCC  1260
             ||||||||||||||||||||||||||||| |||||| |||||||  ||||||||||||||
Sha    1201  TGTTGTTTTTACCTTTCTTGGTAATTTGGATGTCTGTTTTTTAGAAATTAGTTTTCCTCC  1260

Bay-0  1261  TTTTGGTCTTTGTCTCTAGAGATTTTTGTTTATCCGCCGGCTTATGTATGATTTTCGACT  1320
             ||||||||||||||||||||||| ||||||||||||||||||||||||||||||||||||
Sha    1261  TTTTGGTCTTTGTCTCTAGGGATCTTTGTTTATCCGCCGGCTTATGTATGATTTTCGACT  1320

Bay-0  1321  TGTATGTTCTCGTTGCAATTCTAATAAGATAAAAAAAAGCGAGTCACATATCAAAATTTC  1380
             ||||||||||||||||||||||||||||||||||||||||||||||||||||||||||||
Sha    1321  TGTATGTTCTCGTTGCAATTCTAATAAGATAAAAAAAAGCGAGTCACATATCAAAATTTC  1380

Bay-0  1381  AAAAAGCTTGGTACAGTTTTCAAAACATGTAAAACATTTCCAATAACCGTTTTGAAATTC  1440
             ||||||||||||||||||||||||||||||||||||||||||||||||||||||||||||
Sha    1381  AAAAAGCTTGGTACAGTTTTCAAAACATGTAAAACATTTCCAATAACCGTTTTGAAATTC  1440

Bay-0  1441  ATTATTTCAATAACACGGTTTACTTAACCGGTTCTACGTAGTCCACCCTTACCCATTGAC  1500
             ||||||||||||||||||||||||||||||||||||||||||||||||||||||||||||
Sha    1441  GTTATTTCAATAACACGGTTTACTTAACCGGTTCTACGTAGTCCACCCTTACCCATTGAC  1500

Bay-0  1501  ATTTGACACTCTGTCCCGATATAATCTCTTTTGACGAAAGTCAGGCAAAGAAAAAAATGA  1560
             ||||||||||||||||||||||||||||||||||||||||||||||||||||||||||||
Sha    1501  ATTTGACACTCTGTCCCGATATAATCTCTTTTGACGAAAGTCAGGCAAAGAAAAAAATGA  1560

Bay-0  1561  GAATCTTCAAATAAAAAAAAAAGAAGAGACTACGGAGAGTCAAAAACAGTGAGAGCTTTC  1620
             |||||||||||| ||| |||||||||||||||||||||||||||||||||||||||||||
Sha    1561  GAATCTTCAAAT-AAAGAAAAAGAAGAGACTACGGAGAGTCAAAAACAGTGAGAGCTTTC  1620

Bay-0  1621  TTTCATCTTCAGCCAGATCGGAGCAATCTTCTCCATAGATTCCGTTAAACGATGGGAATA  1680
             ||||||||||||||||||||||||||||||||||||||||||||||||||||||||||||
Sha    1621  TTTCATCTTCAGCCAGATCGGAGCAATCTTCTCCATAGATTCCGTTAAACGATGGGAATA  1680

Bay-0  1681  GCCAATAAGCTCCGGCTAAGTTCATCATCTCTCAGCCGAATCCTCCACCGGAGAATACTT  1740
             ||||||||||||||||||||||||||||||||||||||||||||||||||||||||||||
Sha    1681  GCCAATAAGCTCCGGCTAAGTTCATCATCTCTCAGCCGAATCCTCCACCGGAGAATACTT  1740

Bay-0  1741  TACTCATCCGCGGTCAGATCTTTCACCACATCGGAAGGTCACCGTCCCACCATCGTTCAT  1800
             ||||||||||||||||||||||||||||||||||||||||||||||||||||||||||||
Sha    1741  TACTCATCCGCGGTCAGATCTTTCACCACATCGGAAGGTCACCGTCCCACCATCGTTCAT  1800

Bay-0  1801  AAACAAGGTCTCGATATCCTCCATGATCCTTGGTTCAACAAGGTACTTGAAAAACTTGAA  1860
             ||||||||||||||||||||||||||||||||||||||||||||||||||||||||||||
Sha    1801  AAACAAGGTCTCGATATCCTCCATGATCCTTGGTTCAACAAGGTACTTGAAAAACTTGAA  1860

Bay-0  1861  ACTCACTTTCTCAGATTTTGAGTGTGAACACGAACCTGGATCTGATTTTGATTCTTGTTA  1920
             ||||| ||||||||||||||||||||||||||||||||||||||||||||||||||||||
Sha    1861  ACTCAGTTTCTCAGATTTTGAGTGTGAACACGAACCTGGATCTGATTTTGATTCTTGTTA  1920

Bay-0  1921  GGGGACTGCGTTTACGATGACGGAGAGAAATCGTTTAGATCTTAGAGGTCTTCTTCCTCC  1980
             ||||||||||||||||||||||||||||||||||||||||||||||||||||||||||||
Sha    1921  GGGGACTGCGTTTACGATGACGGAGAGAAATCGTTTAGATCTTAGAGGTCTTCTTCCTCC  1980


Bay-0  1981  TAATGTTATGGACTCTGAGCAACAGATTTTTCgttttagtaagttttacttgttgaaact  2040
             ||||||||||||||||||||||||||||||||||||||||||||||||||||||||||||
Sha    1981  TAATGTTATGGACTCTGAGCAACAGATTTTTCGTTTTAGTAAGTTTTACTTGTTGAAACT  2040

Bay-0  2041  ttgtttattgtttatgttgttgtctttagttgttgacatgattgattttgaTATAACAGTG  2100
             ||||||||||||||||||||||||||||||||||||||||||||||||||||||||||||
Sha    2041  TTGTTTATTGTTTATGTTGTTGTCTTAGTTGTTGACATGATTGATTTTGATATAACAGTG  2100

Bay-0  2101  ACTGACCTGAAGAGATTAGAGGAGCAAGCTAGAGATGGACCTTCTGATCCTAATGCTTTG  2160
             ||||||||||||||||| ||||||||||| ||||||||||||||||||||||||||||||
Sha    2101  ACTGACCTGAAGAGATTGGAGGAGCAAGCAAGAGATGGACCTTCTGATCCTAATGCTTTG  2160

Bay-0  2161  GCTAAGTGGCGGATTCTTAATCGGTTACATGACCGAAATGAGACTATGTACTATAAGGTG  2220
             ||||||||||||||||||||||||||||||||||||||||||||||||||||||||||||
Sha    2161  GCTAAGTGGCGGATTCTTAATCGGTTACATGACCGAAATGAGACTATGTACTATAAGGTG  2220

Bay-0  2221  AACTCGTTTTCCTTTTTTGGTCTATTTGCTGTGAAAGATTTTGGTTACTACGCTTGTTTT  2280
             ||||| ||||||||||| ||||||||||||||||||||||||||||||||||||||||||
Sha    2221  AACTCATTTTCCTTTTTCGGTCTATTTGCTGTGAAAGATTTTGGTTACTACGCTTGTTTT  2280

Bay-0  2281  TAGTTTCTGAAGAGGTTATTTACTTCGATTTTTACGTTGCTTTTAGGTTTTGATTAACAA  2340
             ||||||||||||||||||||||||||||||||||||||||||||||||||||||||||||
Sha    2281  TAGTTTCTGAAGAGGTTATTTACTTCGATTTTTACGTTGCTTTTAGGTTTTGATTAACAA  2340

Bay-0  2341  TATTGAGGAGTATGCGCCGATAGTGTATACTCCTACGGTTGGTCTTGTCTGCCAGAACTA  2400
             |||||||||||||||||||||||| |||||||||||||||||||||||||||||||||||
Sha    2341  TATTGAGGAGTATGCGCCGATAGTCTATACTCCTACGGTTGGTCTTGTCTGCCAGAACTA  2400

Bay-0  2401  CAGTGGATTGTTTAGGAGGCCAAGGGGAATGTATTTTAGTGCTGAAGATCGTGGTGAAAT  2460
             ||||||||||||||||||||||||||||||||||||||||||||||||||||||||||||
Sha    2401  CAGTGGATTGTTTAGGAGGCCAAGGGGAATGTATTTTAGTGCTGAAGATCGTGGTGAAAT  2460

Bay-0  2461  GATGTCTATGGTTTACAACTGGCCAGCTGAGCAGGTTCTTCGATATCTCTGACACTGTGC  2520
             ||||||||||||||||||||||||||||||||||||||||||||||||||||||||||||
Sha    2461  GATGTCTATGGTTTACAACTGGCCAGCTGAGCAGGTTCTTCGATATCTCTGACACTGTGC  2520

Bay-0  2521  TTAAGATATCTCAACTCTATTTGGCTGACTAATCCAACGCGTTCTACCCTCTTTCTCTTA  2580
             |||||||||||||||||||||||||||| |||||||||||||||||||||||||||||||
Sha    2521  TTAAGATATCTCAACTCTATTTGGCTGAATAATCCAACGCGTTCTACCCTCTTTCTCTTA  2580

Bay-0  2581  TTGTCTACCAAGGAACTGGTATATCTTTAAGTGACCATTAGTTTTCTGATGTCACGTTTT  2640
             ||||||||||||||||||||||||||||||||||||||||||||||||||||||||||||
Sha    2581  TTGTCTACCAAGGAACTGGTATATCTTTAAGTGACCATTAGTTTTCTGATGTCACGTTTT  2640

Bay-0  2641  TGCTATACATTTAGGTTGATATGATTGTTGTTACCGATGGAAGCCGGATTTTGGGTCTTG  2700
             ||||||||||||||||||||||||||||||||||||||||||||||||||||||||||||
Sha    2641  TGCTATACATTTAGGTTGATATGATTGTTGTTACCGATGGAAGCCGGATTTTGGGTCTTG  2700

Bay-0  2701  GAGATCTAGGTGTTCATGGAATTGGAATTGCTGTAGGGAAGCTTGATTTATATGTTGCCG  2760
             ||||||||||||||||||||||||||||||||||||||||||||||||||||||||||||
Sha    2701  GAGATCTAGGTGTTCATGGAATTGGAATTGCTGTAGGGAAGCTTGATTTATATGTTGCCG  2760

Bay-0  2761  CAGCTGGAATAAATCCTCAACGGGTACTTGCACCACCTAATATTTTCTTATTTTCCCCAT  2820
             ||||||||||||||||||||||||||||||||||||||||||||||||||||||||||||
Sha    2761  CAGCTGGAATAAATCCTCAACGGGTACTTGCACCACCTAATATTTTCTTATTTTCCCCAT  2820

Bay-0  2821  CTCCACTTTTCTGTATGATGTGATACAATACTTATGTGCACATGCTGGTTTAGGTACTAC  2880
             ||||||||||||||||||||||||||||| ||||||||||||||||||||||||||||||
Sha    2821  CTCCACTTTTCTGTATGATGTGATACAATTCTTATGTGCACATGCTGGTTTAGGTACTAC  2880

Bay-0  2881  CTGTCATGATTGATGTGGGAACAAACAATGAAAAACTACGCAATGACCCCATGTGTAAGA  2940
             ||||||||||||||||||||||||||||||||||||||||||||||||||||||||||||
Sha    2881  CTGTCATGATTGATGTGGGAACAAACAATGAAAAACTACGCAATGACCCCATGTGTAAGA  2940


Bay-0  2941  TTCGTTTGCTCCTCTTACCTTCTTAAAGTCAAATTATTGGAGCTGATTATAATAGGTTAA  3000
             ||||||||||||||||||||||||||||||||||||| |||| ||| ||| |||||||||
Sha    2941  TTCGTTTGCTCCTCTTACCTTCTTAAAGTCAAATTATAGGAGATGACTATGATAGGTTAA  3000


Bay-0  3001  TAACGGTTTTCTTCTTTCCGAAAATATATAATTAGTATAACTGTTTACTGAGGTTTGATA  3060
             ||||||||||||||||||||||||||||||||||||||||||||||||||||||||||||
Sha    3001  TAACGGTTTTCTTCTTTCCGAAAATATATAATTAGTATAACTGTTTACTGAGGTTTGATA  3060

Bay-0  3061  TTTTATTATGGTGAACCATATCTATCATCTGTCTCTTTACATAATGAGGTTGAGAATGCA  3120
             ||||||||||||||||||||||||||||||||||||||||||||||||||||||||||||
Sha    3061  TTTTATTATGGTGAACCATATCTATCATCTGTCTCTTTACATAATGAGGTTGAGAATGCA  3120

Bay-0  3121  TATTAtttttttAGTAGGATGGTACAGAATGTGTAGTTGAGATTTGTGTATGCATATATA  3180
             ||||||||| ||||||||||||||||||||||||||||||||||||||||||||||||||
Sha    3121  TATTATTTTCTTAGTAGGATGGTACAGAATGTGTAGTTGAGATTTGTGTATGCATATATA  3180

Bay-0  3181  CATATATTGAAAGGTCAAAGGTTTATAGTATTCTGTATACTGTAGGAGTCTGTCATAAAA  3240
             ||||||||||||||||||||||||||||||||||||||  ||||||||||||||||||||
Sha    3181  CATATATTGAAAGGTCAAAGGTTTATAGTATTCTGTATTATGTAGGAGTCTGTCATAAAA  3240

Bay-0  3241  TTTGGTTGTACGACTTGTTATTTTTGTCTTTCATGGTTTAGGGATTTCCAATAGTTCAGA  3300
             ||||||||||||||||||||||||||||||||||||||||||||||||||||||||||||
Sha    3241  TTTGGTTGTACGACTTGTTATTTTTGTCTTTCATGGTTTAGGGATTTCCAATAGTTCAGA  3300

Bay-0  3301  GACGTGAACGAGAATATCTTATTATCTGTCATGTAATCGTTCTTTCTATGTTTTCTTTTT  3360
             ||| ||||||||||||||||||||||||||||||||||||||||||||||||||||||||
Sha    3301  GACATGAACGAGAATATCTTATTATCTGTCATGTAATCGTTCTTTCTATGTTTTCTTTTT  3360

Bay-0  3361  GACAGACTTGGGTCTGCAACAACGTCGTTTAGAGGATGACGACTATATAGATGTTATCGA  3420
             ||||||||||||||||||||||||||||||||||||||||||||||||||||||||||||
Sha    3361  GACAGACTTGGGTCTGCAACAACGTCGTTTAGAGGATGACGACTATATAGATGTTATCGA  3420

Bay-0  3421  TGAATTTATGGAGGCAGTGTATACTCGGTGGCCACATGTTATTGTGCAGGTAACCGATGC  3480
             ||||||||||||||||||||||||||||||||||||||||||||||||||||||||||||
Sha    3421  TGAATTTATGGAGGCAGTGTATACTCGGTGGCCACATGTTATTGTGCAGGTAACCGATGC  3480

Bay-0  3481  TTTATAGATCTCTGGTATATCAAATGTTTCTTATTATGAAACTTAAATGTTTTTTCTACC  3540
             ||||||||||||||||||||||||||||||||||||||||||||||||||||||||||||
Sha    3481  TTTATAGATCTCTGGTATATCAAATGTTTCTTATTATGAAACTTAAATGTTTTTTCTACC  3540

Bay-0  3541  CACAGTTTGAGGATTTCCAGAGCAAGTGGGCTTTCAAATTATTGCAGAGGTATAGATGCA  3600
             ||||||||||||||||||||||||||||||||||||||||||||||||||||||||||||
Sha    3541  CACAGTTTGAGGATTTCCAGAGCAAGTGGGCTTTCAAATTATTGCAGAGGTATAGATGCA  3600

Bay-0  3601  CCTACCGAATGTTCAATGATGATGTCCAGGTAATGAGTCTTCCGCCTTAGAACGATTCTT  3660
             ||||||||||||||||||||||||||||||||||||||||||||||||||||| ||||||
Sha    3601  CCTACCGAATGTTCAATGATGATGTCCAGGTAATGAGTCTTCCGCCTTAGAACAATTCTT  3660

Bay-0  3661  GTGAAAATTAAATACTAattatttatttatttctgcagtatttttactgtttttagagca  3720
             ||||| ||||||||||||||||||||||||||||||||||||| |||| |||||||||||
Sha    3661  GTGAAGATTAAATACTAATTATTTATTTATTTCTGCAGTATTTATACTATTTTTAGAGCA  3720

Bay-0  3721  atttagtcactactatttatAAAAGTATCATATTTTCTGTGCTCAATCCATCATCTTATC  3780
              |||||||||||||||||||||||||||||||||||||||||||||||||||||||||||
Sha    3721  TTTTAGTCACTACTATTTATAAAAGTATCATATTTTCTGTGCTCAATCCATCATCTTATC  3780

Bay-0  3781  GTTTAATAGTTATTCTTAATCTTTTTTACACGACCAGATGCATTAAAGAGACTGACGAAG  3840
             |||||||||||||||||||| |||||||||||||||||||||||||||||||||||||||
Sha    3781  GTTTAATAGTTATTCTTAATATTTTTTACACGACCAGATGCATTAAAGAGACTGACGAAG  3840

Bay-0  3841  GGAACTTAAAATCTAAATGCTGATACTAGAATCTTGTTTTTCATTCTATGTTTTCATGAT  3900
             ||||||||||||||||||||||||||||||||||||||||||||||||||||||||||||
Sha    3841  GGAACTTAAAATCTAAATGCTGATACTAGAATCTTGTTTTTCATTCTATGTTTTCATGAT  3900

Bay-0  3901  CTCGCTTAGGGGACAGCAGGGGTTGCCATCGCTGGTCTTCTTGGAGCAGTTAGAGCACAG  3960
             |||||||||||||||||||||||||| |||||||||||||||||||||||||||||||||
Sha    3901  CTCGCTTAGGGGACAGCAGGGGTTGCTATCGCTGGTCTTCTTGGAGCAGTTAGAGCACAG  3960


Bay-0  3961  GGGCGACCTATGATTGATTTCCCAAAGATGAAGATCGTTGTTGCTGGCGCGGGAAGGTAA  4020
             ||||||||||||||||||||||||||||||||||||||||||||||||||||||||||||
Sha    3961  GGGCGACCTATGATTGATTTCCCAAAGATGAAGATCGTTGTTGCTGGCGCGGGAAGGTAA  4020

Bay-0  4021  GCCATTCTCGGTTGGTGTTTTTCTCGTCTTTCCTGGTCCTAGTTTACTAGAAATTGGTAT  4080
             ||||||||||||||||||||||||||||||||||||||||||||||||||||||||||||
Sha    4021  GCCATTCTCGGTTGGTGTTTTTCTCGTCTTTCCTGGTCCTAGTTTACTAGAAATTGGTAT  4080

Bay-0  4081  TGGTAAAAGATTGGTCTTGCTATTACTGTTTAATTATTTGTTGGTCAGGGATCTTTCGTT  4140
             |||||||||||||||||||||||||||||||| |||||| ||||||||||||||||||||
Sha    4081  TGGTAAAAGATTGGTCTTGCTATTACTGTTTAGTTATTTATTGGTCAGGGATCTTTCGTT  4140

Bay-0  4141  TATTTGTTGGTTTTGAGTAATGTTTGAAGAGATTTAAAATGAAAACTTTCAGCTGCATCA  4200
             |||||||| |||||||||||||||||||||||||||||||||||||||||||||||||||
Sha    4141  TATTTGTTAGTTTTGAGTAATGTTTGAAGAGATTTAAAATGAAAACTTTCAGCTGCATCA  4200

Bay-0  4201  CTGCATAAGTTAGCTTAGACTAGTTATATTAGAATACTGAGCTTCCTTGTAAAAATCTTT  4260
             ||||||||||||||||||||||||||||||||||||||||||||||||||||||||||||
Sha    4201  CTGCATAAGTTAGCTTAGACTAGTTATATTAGAATACTGAGCTTCCTTGTAAAAATCTTT  4260

Bay-0  4261  GATTTCTGCAGTGCGGGAATTGGTGTTCTTAATGCTGCGAGGAAGACAATGGCACGAATG  4320
             ||||||||||||||||||||||||||||||||||||||||||||||||||||||||||||
Sha    4261  GATTTCTGCAGTGCGGGAATTGGTGTTCTTAATGCTGCGAGGAAGACAATGGCACGAATG  4320

Bay-0  4321  TTGGGAAATACTGAAACTGCATTTGATAGTGCACAAAGTCAATTTTGGGTGGTTGATGCG  4380
             ||||||||||||||||||||||||||||||||||||||||||||||||||||||||||||
Sha    4321  TTGGGAAATACTGAAACTGCATTTGATAGTGCACAAAGTCAATTTTGGGTGGTTGATGCG  4380

Bay-0  4381  CAGGTATGTTATTCTATAGTGGAGTAATTGATTTTCAGATTAGGGAGCTGATAAGACGTT  4440
             ||||||||||||| ||||||||||||||||||||||||||||||||||||||||||||||
Sha    4381  CAGGTATGTTATTGTATAGTGGAGTAATTGATTTTCAGATTAGGGAGCTGATAAGACGTT  4440

Bay-0  4441  TTAGCCAGTCAAGCTATTTCCCTCTGGTTTCCTTTTACTGCTGACGAAATCACTTCATCA  4500
             ||||||||||||||||||||||||||||||||||||||||||||||||||||||||||||
Sha    4441  TTAGCCAGTCAAGCTATTTCCCTCTGGTTTCCTTTTACTGCTGACGAAATCACTTCATCA  4500

Bay-0  4501  TCAttttttttttGGGCGTGTGTTTTGGTCCAATAGCTTAAGGAGCTGAAGTTGTGAACT  4560
             |||||||||||||||||||||||||||||||||||||| |||||||||||||||||||||
Sha    4501  TCATTTTTTTTTTGGGCGTGTGTTTTGGTCCAATAGCTCAAGGAGCTGAAGTTGTGAACT  4560

Bay-0  4561  CttttttttGCACTTAAATCAGGGTCTTATCACCGAAGGCCGGGAAAATATTGACCCAGA  4620
             ||||||||||||||||||||||||||||||||||||||||||||||||||||||||||||
Sha    4561  CTTTTTTTTGCACTTAAATCAGGGTCTTATCACCGAAGGCCGGGAAAATATTGACCCAGA  4620

Bay-0  4621  GGCTCAGCCTTTTGCTAGGAAGACTAAAGAAATGGAGCGTCAGGGATTAAAAGAAGGAGC  4680
             ||||||||||||||||||||||||||||||||||||||||||||||||||||||||||||
Sha    4621  GGCTCAGCCTTTTGCTAGGAAGACTAAAGAAATGGAGCGTCAGGGATTAAAAGAAGGAGC  4680

Bay-0  4681  AACTCTTGTGGAAGTGGTCTGTTATTCTTTCTATCAATATGTTCATCAAGTATACTAAAT  4740
             ||||||||||||||||||||||||||||||||||||||||||||||||||||||||||||
Sha    4681  AACTCTTGTGGAAGTGGTCTGTTATTCTTTCTATCAATATGTTCATCAAGTATACTAAAT  4740

Bay-0  4741  CTAATTTGGCAATGAGTCTGGTGTTGGAGAGAAATTCTCCCTTAAAAAAGTTTCCCTGCT  4800
             |||||||||||||||| |||||||||||||||||||||||||||||||||||||||||||
Sha    4741  CTAATTTGGCAATGAGCCTGGTGTTGGAGAGAAATTCTCCCTTAAAAAAGTTTCCCTGCT  4800

Bay-0  4801  TAATACTGACAAGAAACAACGGTTTTCTGTTTCATCTGTGCTAGGTTCGTGAAGTTAAAC  4860
             ||||||||||||||||||||||||||||||||||||||||||||||||||||||||||||
Sha    4801  TAATACTGACAAGAAACAACGGTTTTCTGTTTCATCTGTGCTAGGTTCGTGAAGTTAAAC  4860

Bay-0  4861  CTGATGTGCTTCTTGGTTTATCAGCAGTTGGAGGGTTATTCTCAAAAGAGGTAATGAAAG  4920
             ||||||||||||||||||||||||||||||||||||||||||||||||||||||||||||
Sha    4861  CTGATGTGCTTCTTGGTTTATCAGCAGTTGGAGGGTTATTCTCAAAAGAGGTAATGAAAG  4920

Bay-0  4921  CTGCTAAATGGCTAAACATATCTAGTGCTATGCAACAGAATCAGTGATTTAGCTAATTAG  4980
             |||| |||||||||||||||||||||||||||||||||||||||||||||||||||||||
Sha    4921  CTGCCAAATGGCTAAACATATCTAGTGCTATGCAACAGAATCAGTGATTTAGCTAATTAG  4980

Bay-0  4981  ATGAGCGCGCAAATGAAATCTAGTATTTCTTCAGTGACCACTGTCTTCATAATCTGGTTT  5040
             ||||||||||||||||||||||||||||||||||||||||||||||||||||||||||||
Sha    4981  ATGAGCGCGCAAATGAAATCTAGTATTTCTTCAGTGACCACTGTCTTCATAATCTGGTTT  5040

Bay-0  5041  CCATGGTAACTGCTAAGTAGATGAGTGAGCTAGTAGAATGTAGCATAGCTCATTGCTTGC  5100
             ||||||||||||||||||||||||||||||||||||||||||||||||||||||||||||
Sha    5041  CCATGGTAACTGCTAAGTAGATGAGTGAGCTAGTAGAATGTAGCATAGCTCATTGCTTGC  5100

Bay-0  5101  ATGATTCTGACATATGGTAGTTGATTTTTGAAGTCATTGTTGGGATATTGTGTTTAAGGT  5160
             ||||||||||||||||||||||||| ||||||||||||||||||||||||||||||||||
Sha    5101  ATGATTCTGACATATGGTAGTTGATATTTGAAGTCATTGTTGGGATATTGTGTTTAAGGT  5160

Bay-0  5161  TTTAGAAGCAATGAAAGGTTCAACCTCGACAAGGCCTGCTATATTTGCAATGTCAAACCC  5220
             ||||||||||||||||||||||||||||||||||||||||||||||||||||||||||||
Sha    5161  TTTAGAAGCAATGAAAGGTTCAACCTCGACAAGGCCTGCTATATTTGCAATGTCAAACCC  5220

Bay-0  5221  CACAAAAAATGGTAATTTGTCTTCTTTCTTTGCCATAAAATGTGTTTTGCGGTTATTTGT  5280
             ||||||||||||||||||||||||||||||||||||||||||||||||||||||||||||
Sha    5221  CACAAAAAATGGTA-TTTGTCTTCTTTCTTTGCCATAAAATGTGTTTTGCGGTTATTTGT  5280

Bay-0  5281  GATTCTTTTTTGGATAGATGTAATAGTTACTTGTGTTTTGAATAACAGCTGAATGCACGC  5340
             |||||||||||||||||||||||||| |||||||||||||||||||||||||||||||||
Sha    5281  GATTCTTTTTTGGATAGATGTAATAGGTACTTGTGTTTTGAATAACAGCTGAATGCACGC  5340

Bay-0  5341  CTCAAGATGCATTTTCGATATTAGGCGAAAATATGATTTTTGCGAGTGGAAGCCCATTCA  5400
             |||||||||| |||||||||||||||||||||||||||||||||||||||||||||||||
Sha    5341  CTCAAGATGCTTTTTCGATATTAGGCGAAAATATGATTTTTGCGAGTGGAAGCCCATTCA  5400

Bay-0  5401  AAAATGTGGAATTTGGTATGTATGAGGACCTCTGTATCATTTAGCATACGCCATTTTATA  5460
             ||||||||||||||||||||||||||||||||||||||||||||||||||||||||||||
Sha    5401  AAAATGTGGAATTTGGTATGTATGAGGACCTCTGTATCATTTAGCATACGCCATTTTATA  5460

Bay-0  5461  TGAGGCTTTTTCAAATATTAGTACCTTTTCTTGTGGCAGGAAATGGTCATGTAGGACATT  5520
             ||||||||||||||||||||||||||||||||||||||||||||||||||||||||||||
Sha    5461  TGAGGCTTTTTCAAATATTAGTACCTTTTCTTGTGGCAGGAAATGGTCATGTAGGACATT  5520

Bay-0  5521  GCAACCAGGGAAACAACATGTATCTATTTCCAGGGTTGGTTCCTCTTGTTTATTGTCCGA  5580
             ||||||||||||||||||||||||||||||||||||||||||||||||||||||||||||
Sha    5521  GCAACCAGGGAAACAACATGTATCTATTTCCAGGGTTGGTTCCTCTTGTTTATTGTCCGA  5580

Bay-0  5581  AAGAATTGCATACATTCTTGAATTGATACATTAATTGTCATGAAATAAATTGATTTTCTA  5640
             ||||||||||||||||||||||||||||||||||||||||||||||||||||||||||||
Sha    5581  AAGAATTGCATACATTCTTGAATTGATACATTAATTGTCATGAAATAAATTGATTTTCTA  5640

Bay-0  5641  ATCCTGTAAAATTTATTGTAGTATTGGACTTGGCACTCTTCTATCTGGTGCTCCCATTGT  5700
             ||||||||||||||| ||||||||||||||||||||||||||||||||||||||||||||
Sha    5641  ATCCTGTAAAATTTAATGTAGTATTGGACTTGGCACTCTTCTATCTGGTGCTCCCATTGT  5700

Bay-0  5701  CTCCGACGGGATGCTTCAGGCCGCATCCGAGTGGTAAGCTCCTTTCTTTACCTTCTGCAT  5760
             |||||||||||||||||||||||||||||||||||||||||||||||||||||| |||||
Sha    5701  CTCCGACGGGATGCTTCAGGCCGCATCCGAGTGGTAAGCTCCTTTCTTTACCTTTTGCAT  5760

Bay-0  5761  TTATCATTCCTGATTTATTCTCAACTAAAGCTGAAGTGAATACCATGATATATAACTACG  5820
             ||||||||||||||||||||||||||||||||||||||||||||||||||||||||||||
Sha    5761  TTATCATTCCTGATTTATTCTCAACTAAAGCTGAAGTGAATACCATGATATATAACTACG  5820

Bay-0  5821  GACCAAATTTAAATGATTATTAAGAATGTCCGCATATTTGGCTGATGTCACGAACTATAA  5880
             ||||||||||||||||||||||||||||||||||||||||||||||||||||||||||||
Sha    5821  GACCAAATTTAAATGATTATTAAGAATGTCCGCATATTTGGCTGATGTCACGAACTATAA  5880

Bay-0  5881  CATACCCCCCATCCACGAGACAGGAACCACAGTAGTTCGACATGTTCTCTCCAAATGGTA  5940
             ||||||||||||||||||||||||||||||||||||||||||||||||||||||||||||
Sha    5881  CATACCCCCCATCCACGAGACAGGAACCACAGTAGTTCGACATGTTCTCTCCAAATGGTA  5940


Bay-0  5941  TTCCCAAAAGTAGGCCTTCGGTTGAGGTTTCATATTGTAGTTAATGTAAAACTCATATTT  6000
             ||||||||||||||||||||||||||||||||||||||||||||||||||||||||||||
Sha    5941  TTCCCAAAAGTAGGCCTTCGGTTGAGGTTTCATATTGTAGTTAATGTAAAACTCATATTT  6000

Bay-0  6001  TTAAGGAGACCATGAAAATAAACAAGTCCCGTAATTTATGAATGTGGAATCATGACATTG  6060
             ||||||||||||||||||||||||||||||||||||||||||||||||||||||||||||
Sha    6001  TTAAGGAGACCATGAAAATAAACAAGTCCCGTAATTTATGAATGTGGAATCATGACATTG  6060

Bay-0  6061  TTCTGCTAAACTACTGCAGCCTAGCAGCATACATGAGCGAAGAAGAAGTGCTAGAGGGGA  6120
             ||||||||||| ||||||||||||||||||||||||||||||||||||||||||||||||
Sha    6061  TTCTGCTAAACAACTGCAGCCTAGCAGCATACATGAGCGAAGAAGAAGTGCTAGAGGGGA  6120

Bay-0  6121  TTATATACCCTCCAATATCAAGGTAACAGTAACACAATCAAAGAACATCCTTACTGTTAC  6180
             ||||||||||||||||||||||||||||||||||||||||||||||||||||||||||||
Sha    6121  TTATATACCCTCCAATATCAAGGTAACAGTAACACAATCAAAGAACATCCTTACTGTTAC  6180

Bay-0  6181  ATAAGAAAATGTTTTGATATAGTTGCTACATTTGGCATGTGGAATTGAAATTTCAGGATA  6240
             ||||||||||||||||||||||||||||||||||||||||||||||||||||||||||||
Sha    6181  ATAAGAAAATGTTTTGATATAGTTGCTACATTTGGCATGTGGAATTGAAATTTCAGGATA  6240

Bay-0  6241  CGAGACATAACAAAAAGAATAGCAGCTGCGGTGATTAAGGAAGCGATTGAGGAGGATTTG  6300
             ||||||||||||||||||||||||||||||||||||||||||||||||||||||||||||
Sha    6241  CGAGACATAACAAAAAGAATAGCAGCTGCGGTGATTAAGGAAGCGATTGAGGAGGATTTG  6300

Bay-0  6301  GTTGAAGGATATAGAGAAATGGACGCTCGAGAGATCCAAAAGCTGGATGAGGTAACAATT  6360
             ||||||||||||||||||||||||||||||||||||||||||||||||||||||||||||
Sha    6301  GTTGAAGGATATAGAGAAATGGACGCTCGAGAGATCCAAAAGCTGGATGAGGTAACAATT  6360

Bay-0  6361  CAGCTAATGACTGGCTTTAATATAATCATAAATGATAGAAAAAGAAAAGATGTCAAAACA  6420
             |||||||||||| |||||||||||||||||||||||||||||||||||||||||||||||
Sha    6361  CAGCTAATGACTTGCTTTAATATAATCATAAATGATAGAAAAAGAAAAGATGTCAAAACA  6420

Bay-0  6421  ACATCCCATCTTTCACGTGTTTTGGACTCCTGATCATAAAATCGTATTCTACGAAGAATG  6480
             ||||| ||||||||||||||||||||||||||||||||||||||||||||||||||||||
Sha    6421  ACATCGCATCTTTCACGTGTTTTGGACTCCTGATCATAAAATCGTATTCTACGAAGAATG  6480

Bay-0  6481  TCGATCGGGTTTGATTATATGTAAAGAACAGAAGAGTGTTTACCTGATTTCTCACTTAGA  6540
             ||||||||||||||||||||||||||||||||||||||||||||||||||||||||||||
Sha    6481  TCGATCGGGTTTGATTATATGTAAAGAACAGAAGAGTGTTTACCTGATTTCTCACTTAGA  6540

Bay-0  6541  GAAAATAAAGACTAAACCAGAAACTAAGCCTAAATGTTGGTTATGTTGACGCGGTCTTGT  6600
             |||||||||||||||||||||||||||||| ||||||| |||||||||||||||||||||
Sha    6541  GAAAATAAAGACTAAACCAGAAACTAAGCCGAAATGTTAGTTATGTTGACGCGGTCTTGT  6600

Bay-0  6601  GTCTTCCGCAGGAGGGGCTGATGGAGTATGTGGAAAACAACATGTGGAATCCAGAGTACC  6660
             ||||||||||||||||||||||||||||||||||||||||||||||||||||||||||||
Sha    6601  GTCTTCCGCAGGAGGGGCTGATGGAGTATGTGGAAAACAACATGTGGAATCCAGAGTACC  6660

Bay-0  6661  CGACTTTGGTCTACAAGGATGACTAAGTTTGGTTCCCAAAATGAATAAGTCTACTTTCTC  6720
             ||||||||||||||||||||||||||||||||||||||||||||||||||||||||||||
Sha    6661  CGACTTTGGTCTACAAGGATGACTAAGTTTGGTTCCCAAAATGAATAAGTCTACTTTCTC  6720

Bay-0  6721  TTATTTATATGTAAAAATTTATAATTGGGGTTGCTTCATACCCAATCAAATTCATAGCTA  6780
             ||||||||||||||||||||||||||||||||||||||||||||||||||||||||||||
Sha    6721  TTATTTATATGTAAAAATTTATAATTGGGGTTGCTTCATACCCAATCAAATTCATAGCTA  6780

Bay-0  6781  CTGCTTTCTCTTGTTCTTATGTaaaaaaaaaaaaaaTATATAATTGGGTTGTTTGTTATC  6840
             ||||||||||||||||||||||||||||||||||||||||||||||||||||||||||||
Sha    6781  CTGCTTTCTCTTGTTCTTATGTAAAAAAAAAAAAAATATATAATTGGGTTGTTTGTTATC  6840

Bay-0  6841  CAATCAGATTCATAGCTTCCACTTTCTCTTATTTATATGTAAAAAAGAAGTTATATTCAA  6900
             |||||||| |||||||||||||||||||||||||||||||||||||||||||||||||||
Sha    6841  CAATCAGAGTCATAGCTTCCACTTTCTCTTATTTATATGTAAAAAAGAAGTTATATTCAA  6900

Bay-0  6901  TCAGATTCATAGCAGCAACATTTTACATACATCGTATTAGAAATGCATTTTCTTTTATAA  6960
             ||||||||||||||||||||||||||||||||||||||||||||||||||||||||||||
Sha    6901  TCAGATTCATAGCAGCAACATTTTACATACATCGTATTAGAAATGCATTTTCTTTTATAA  6960

Bay-0  6961  CTTATAGTTTCAGGTCATGATACTAAGAGTAGTGAGTGAGGAACAATTTGAAGTTCTTTA  7020
             ||||||||||||||||||||||||||||||||||||||||||||||||||||||||||||
Sha    6961  CTTATAGTTTCAGGTCATGATACTAAGAGTAGTGAGTGAGGAACAATTTGAAGTTCTTTA  7020

Bay-0  7021  ATTCAAGCAATCATCTTTATCAATCATAATATGAGTAAATGAATGAGAAACAATACACCA  7080
             ||||||||||||||||||||||||||||||||||||||||||||||||||||||||||||
Sha    7021  ATTCAAGCAATCATCTTTATCAATCATAATATGAGTAAATGAATGAGAAACAATACACCA  7080

Bay-0  7081  TTCTCTTCCAATTAAGCTATTCTCTACTCGAAACCTTATAAGCTAGACCACTAACTCTCA  7140
             ||||||||||||||||||||||||||||||||||||||||||||||||||||||||||||
Sha    7081  TTCTCTTCCAATTAAGCTATTCTCTACTCGAAACCTTATAAGCTAGACCACTAACTCTCA  7140

Bay-0  7141  AACCGTAGGTGGTAAGTATTCTAATCCAATTAGCAGCATGGAATTAGTTTCTACTGTTGT  7200
             ||||||||||||||||||||||||||||||||||||||||||||||||||||||||||||
Sha    7141  AACCGTAGGTGGTAAGTATTCTAATCCAATTAGCAGCATGGAATTAGTTTCTACTGTTGT  7200

Bay-0  7201  AAATCATGAGAGATTGCTATATTTGCTCGAGACGAACTAATCCGGAAAATATATGAAGAC  7260
             ||||||||||||||||||||||||||||||||||||||||||||||||||||||||||||
Sha    7201  AAATCATGAGAGATTGCTATATTTGCTCGAGACGAACTAATCCGGAAAATATATGAAGAC  7260

Bay-0  7261  GAATTTCTGTTACACAAAACCATGGGTATACTGAAAATATCAAAATGTACGTGTCTAGTG  7320
             ||||||||||||||||||||||||||||||||||||||||||||||||||||||||||||
Sha    7261  GAATTTCTGTTACACAAAACCATGGGTATACTGAAAATATCAAAATGTACGTGTCTAGTG  7320
```

**Figure S2.** Alignment of genomic DNA including promoter, ORF (green) and introns of NAD-ME1 from Bay-0 and Sha. SNPs between genotypes are highlighted in red.

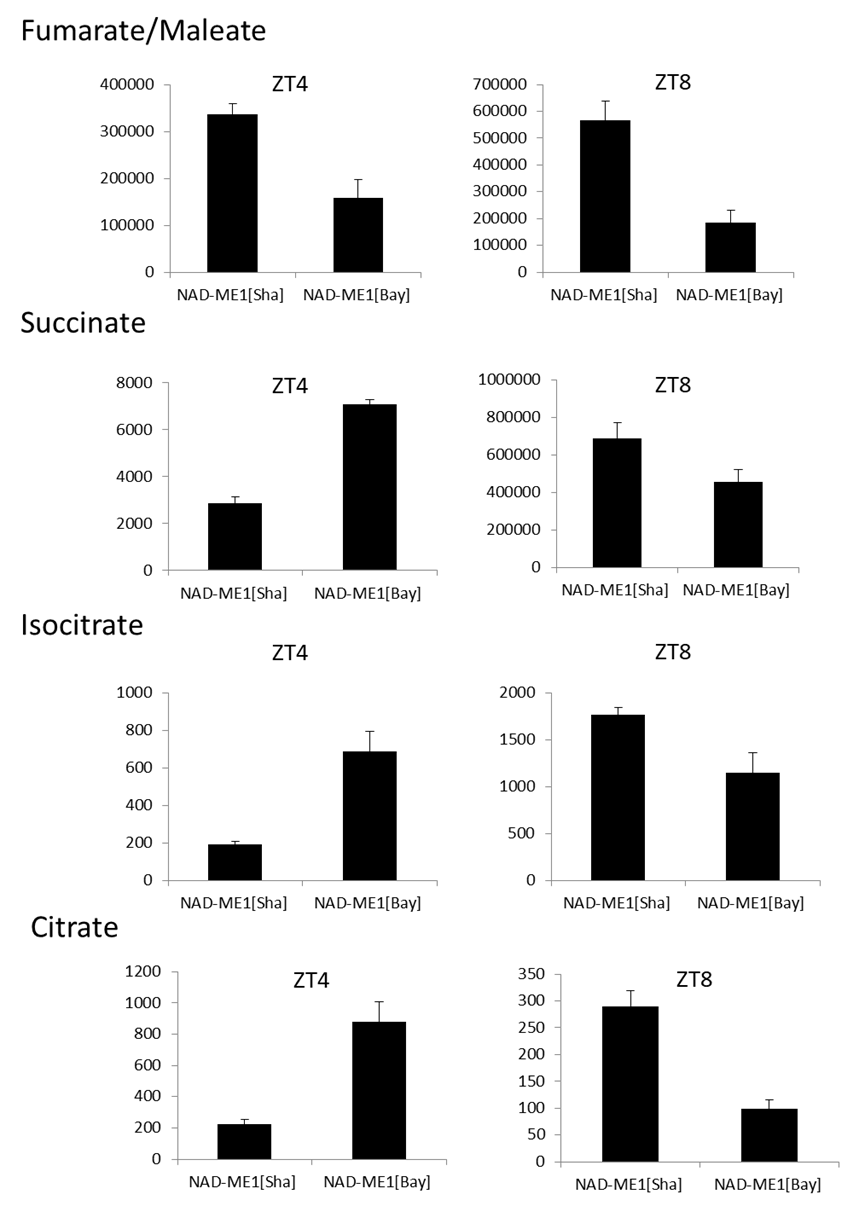

**a**

**b**

**b**

**b**

**b**

**b**

**b**

**b**

**b**

**a**

**a**

**a**

**a**

**a**

**a**

**a**

**Figure S3. Impact of NAD-ME1 alleles in Arabidopsis TCA metabolites.** Ion intensity of fumarate/maleate, succinate, isocitrate and citrate analyzed by UPLC-Q-TOF-MS/MS from NAD-ME[Sha] and NAD-ME1[Bay-0] leaves at two different harvest times, ZT4 and ZT8. Different letters indicate statistically significant differences determined by analysis of variance (ANOVA) with post‐hoc Tukey’s honestly significant difference (HSD) test (P < 0.05). Error lines represent ± standard deviation of the mean.
